# Attenuating midline thalamus bursting to mitigate absence epilepsy

**DOI:** 10.1101/2023.09.18.558258

**Authors:** Ping Dong, Konstantin Bakhurin, Yuhui Li, Mohamad A. Mikati, Jianmin Cui, Warren M. Grill, Henry H. Yin, Huanghe Yang

**Affiliations:** Department of Biochemistry, Duke University Medical Center, Durham, NC 27710, USA; Department of Psychology and Neuroscience, Duke University, Durham, NC 27708, USA; Department of Biomedical Engineering, Duke University, Durham, NC, 27708, USA; Department of Neurobiology, Duke University Medical Center, Durham, NC 27710, USA; Department of Pediatrics, Duke University Medical Center, Durham, NC 27710, USA; Department of Biomedical Engineering, Washington University in St. Louis, MO 63130, USA; Department of Neurosurgery, Duke University Medical Center, Durham, NC 27710, USA

## Abstract

Advancing the mechanistic understanding of absence epilepsy is crucial for developing new therapeutics, especially for patients unresponsive to current treatments. Utilizing a recently developed mouse model of absence epilepsy carrying the BK gain-of-function channelopathy D434G, here we report that attenuating the burst firing of midline thalamus (MLT) neurons effectively prevents absence seizures. We found that enhanced BK channel activity in the BK-D434G MLT neurons promotes synchronized bursting during the ictal phase of absence seizures. Modulating MLT neurons through pharmacological reagents, optogenetic stimulation, or deep brain stimulation effectively attenuates burst firing, leading to reduced absence seizure frequency and increased vigilance. Additionally, enhancing vigilance by amphetamine, a stimulant medication, or physical perturbation also effectively suppresses MLT bursting and prevents absence seizures. These findings suggest that the MLT is a promising target for clinical interventions. Our diverse approaches offer valuable insights for developing new therapeutics to treat absence epilepsy.

**Highlights:** The midline thalamus (MLT) is a key thalamic region for absence seizure pathogenesis MLT neurons exhibit synchronized bursting during ictal phase. BK channel contributes to MLT burst firing Attenuating MLT bursting increases vigilance and suppresses absence seizures

## Introduction

Pediatric absence epilepsy affects 2-8 out of every 100,000 children worldwide^1–3^. It is characterized by brief lapses of consciousness and reduction of vigilance accompanied by generalized, synchronous, bilateral 2.5-4 Hz spike and slow-wave discharges (SWDs) recorded by electroencephalogram (EEG)^2–9^. Despite the availability of anti-absence medications such as ethosuximide (ESM), valproic acid and lamotrigine, about 30% of the patients remain pharmaco-resistant and face long-term neuropsychiatric comorbidities, including attention, mood, cognitive, and memory impairments^2–6^. This underscores the importance and urgency of gaining a deeper understanding of the pathogenic mechanisms of absence epilepsy, identifying new therapeutic targets, and developing more effective treatments.

The established findings from earlier clinical and animal research indicate that rhythmic oscillations in the cortical-thalamic reticular nucleus (TRN)-thalamic system are crucial for producing and transmitting SWDs^2, 5, 8^. The abnormal rhythmic oscillations lead to hypersynchronization within this circuit, resulting in SWDs and a transient loss of consciousness or reduction of vigilance^7^. The hyperactivity in the glutamatergic thalamocortical (TC) neurons and the GABAergic TRN neurons, driven by bursts of action potentials via T-type voltage-gated Ca^2+^ channels (TCaVs) during every SWD cycle, underpin the rhythm-generating mechanism of absence seizures^3, 8, 10^. In this hypersynchronization state, multiple TRN cells fire simultaneously, producing strong IPSPs onto TC cells. Due to high levels of T-CaV channels in thalamocortical cells, this strong inhibition results in postinhibitory rebound bursts that re-engage both TRN and cortex^5^.

Targeting rhythmic oscillations to prevent hypersynchronization is therefore essential for treating absence seizures. As burst firing of thalamic neurons is important in driving the rhythmic oscillations and hypersynchronization^11^, targeting thalamic bursting could be a promising strategy to treat absence epilepsy. However, the thalamus consists of distinct sub-nuclei that vary in size, function, and connectivity^12, 13^. Therefore, accurate identification and precise targeting of the thalamic regions that are important for absence seizure pathogenesis are crucial for the development of innovative therapeutic strategies to effectively treat absence epilepsy.

We recently reported a new mouse model for absence epilepsy with direct human relevance^14^. The mouse model carries the gain-of-function (GOF) D434G variant of Ca^2+^-activated, large-conductance BK K^+^ channel, which was identified from a large family of patients with generalized absence seizures and/or paroxysmal dyskinesia^15, 16^. The BK-D434G mice recapitulate human manifestations of absence seizures, including frequent SWDs accompanied by behavioral arrest, reduced vigilance, and a positive response to first-line anti-absence medications, ESM and valproate^14^.

Utilizing the new absence seizure mouse model with direct human relevance, here we report that hypersynchronized bursting of the midline thalamus (MLT) plays a critical role for absence seizure pathogenies. Pharmacological inhibition, optogenetic stimulation or electrical neuromodulation of the BK-D434G MLT effectively suppresses its bursting, enhances vigilance and alleviates absence seizures. Furthermore, enhancing vigilance with amphetamine or physical perturbation also suppresses MLT bursting and prevents absence seizures. Our study thus identifies the MLT as a key thalamic region for targeted anti-absence interventions, uncovers MLT hypersynchronized burst firing in absence seizure pathogenesis, discovers the role of BK channels in promoting burst firing of MLT neurons, and provides a variety of approaches to attenuate MLT bursting, which have the potential to be translated into novel treatments of absence epilepsy of different etiologies.

## Results

### Absence epilepsy induce c-Fos expression in BK-D434G midline thalamus

Homozygous BK-D434G (BK^DG/DG^) mice develop frequent absence seizures (> 250 SWD per hour) with concomitant behavioral arrest^14^. To identify the key loci for absence seizure pathogenesis, we explored the cortical-TRN-thalamus circuit, known to be responsible for absence seizure genesis^2, 3, 5^. We utilized c-Fos, an immediate-early gene that reports both increases in firing rate and changes in firing pattern^17–22^. We observe strong c-Fos staining in the cortex and the MLT, especially in the intermediodorsal (IMD) and central medial (CM) nuclei in BK^DG/DG^ mice but not in wildtype (BK^WT/WT^) control mice (**Figs. 1A-C, S1-C**). Interestingly, no obvious c-Fos staining was observed in the BK^DG/DG^ TRN or ventral posteromedial nucleus (VPM) (**Fig. S1D**), two thalamic regions that were implicated in absence seizure pathogenesis of the spontaneous mutation rodent models, such as GAERS rats^23, 24^.

**Figure 1.**
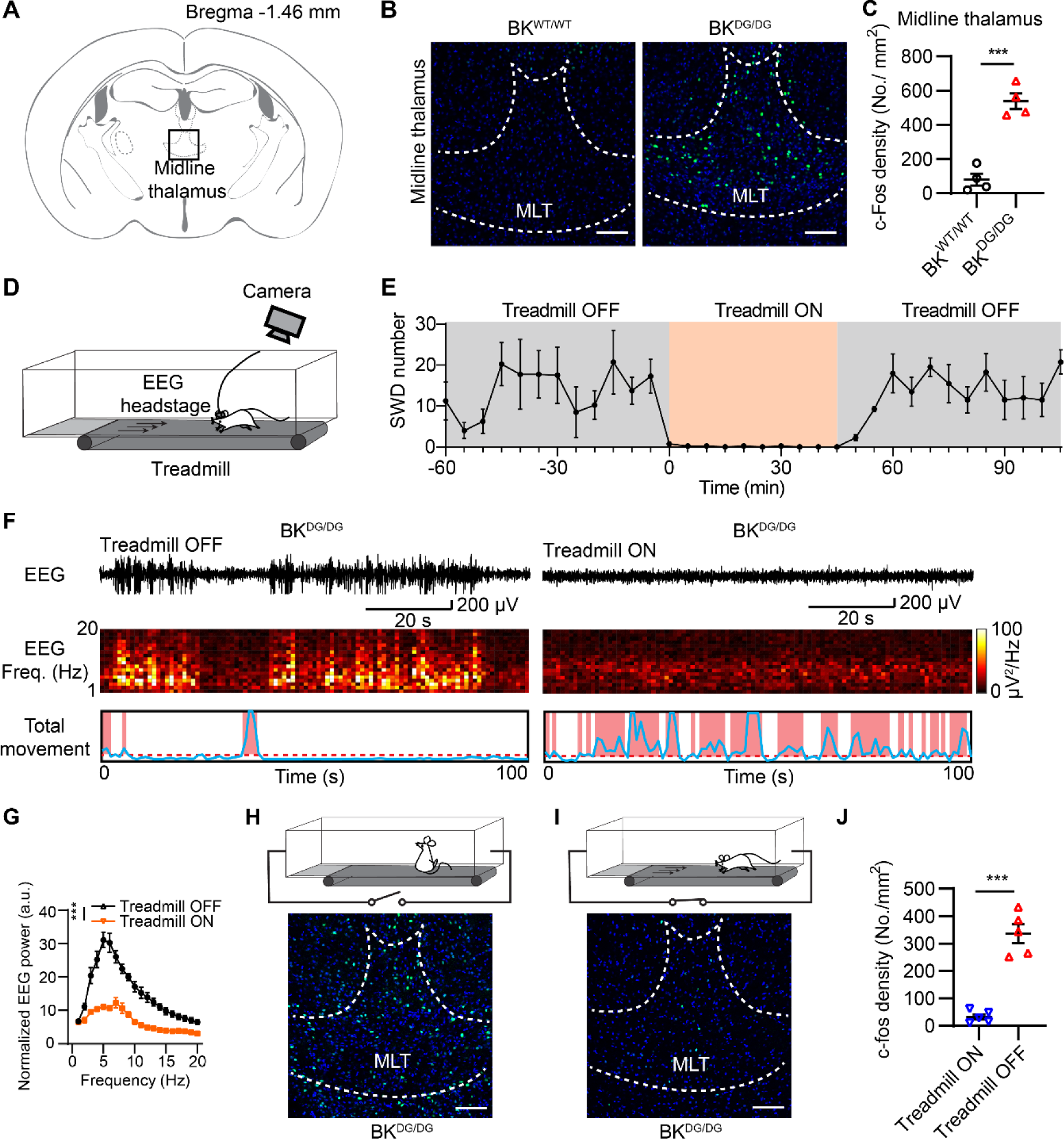
c-Fos expression in BK-D434G midline thalamic (MLT) neurons is positively correlated with absence seizures but inversely correlated with vigilance level. (A) A coronal atlas showing the location of MLT in the brain. (B) Representative images of c-Fos expression in the MLT of the *BK^WT/WT^* (left) and *BK^DG/DG^* (right) mice. Scale bar 100 μm. (C) Statistics of c-Fos density in the MLT (Two-tailed unpaired Student’s *t*-test, t_6_ = 7.944, *P* = 0.0002). n = 4 mice per group. *** *P* < 0.001. (D) Schematic of the simultaneous video-electroencephalogram (EEG) recording of freely moving mice on treadmill. (E) Time course of the treadmill effect on the spontaneous SWDs of the *BK^DG/DG^* mice. The orange box indicates the period when the treadmill was on. (F) Representative EEG traces, corresponding spectrograms, and total movement from a *BK^DG/DG^* mouse before (Left panel, Treadmill OFF) and during treadmill (Right panel, Treadmill ON). The treadmill was set at lowest speed (2 m/min) to avoid fatigue. (G) Comparison of the power spectral density of EEG recorded from *BK^DG/DG^*mice during treadmill OFF and treadmill ON. Two-way ANOVA, *F*_(1,6)_ = 122.3, *P* < 0.0001. n = 4 mice. (H, I) Schematic drawing of the experimental design (top) and representative images of c-Fos expression (bottom) in the MLT from *BK^DG/DG^* under two conditions: when the animals were habituated on a stationary treadmill (H) or on a moving treadmill (I). (J) Treadmill effect on c-Fos density in the BK-D434G MLT. Two-tailed unpaired Student’s *t*-test, t_8_ = 8.418, *P* < 0.0001. n = 5 mice per group. In all plots and statistical tests, summary graphs show mean ± s.e.m.

The MLT is an important brain region in regulating vigilance and arousal^25–28^. Intriguingly, absence seizures in patients typically occur during the drowsy or passive awake state, when their vigilance level decreases^29^. Conversely, when their vigilance level is sustained through engaging with a new toy or participating in a more engaging task, the incidence of absence seizures dramatically diminishes^30, 31^. To test whether the upregulation of c-Fos expression in the BK-D434G MLT is correlated with the vigilance level of the animals, we maintained their vigilance level by constantly stimulating the BK^DG/DG^ mice with a low speed treadmill equipped with a video-EEG recording system. This setup allowed us to manipulate vigilance levels by switching the treadmill on or off (**Fig. 1D**). After habituating in the treadmill chamber for one hour, the mice developed frequent absence seizures accompanied with behavioral arrest (**Fig. 1E-F**). Once the treadmill was turned on, the animals were forced to move, and their spontaneous SWDs immediately disappeared. After the treadmill was turned off, spontaneous SWDs gradually appeared again (**Fig. 1E**). In addition, the c-Fos expression in the MLT was significantly reduced in the ‘treadmill ON’ group compared with the control ‘treadmill OFF’ group (**Fig. 1 H-J**). All these findings suggest that c-Fos expression in BK-D434G MLT neurons is positively correlated with absence seizures but inversely correlated with vigilance level. The MLT thus could be an important thalamic region for the absence seizure pathophysiology of BK-D434G mice.

### BK-D434G MLT shows hypersynchronized burst firing during ictal phase *in vivo*

To understand the contribution of the MLT to absence seizure pathogenesis, we conducted *in vivo* electrophysiological recording of freely behaving BK^DG/DG^ mice with simultaneous EEG recording and single-unit recording from MLT neurons (**Figs. 2A-B, S2-B**). After aligning single-unit (**Fig. 2C**) and EEG signals to the SWD positive peaks (**Figs. 2B** and **S3**, see Methods), we observed that BK^DG/DG^ MLT neurons exhibit a significant increase of burst firing (> 150 Hz) at the SWD positive peaks (**Fig. 2D-E**). By comparing the MLT neuronal spikes during inter-ictal (non-SWD) and ictal (SWD) phases, we found that the averaged BK^DG/DG^ MLT firing rate was reduced during the ictal phase (**Fig. 2F-G**). In addition, MLT firing switched from unsynchronized activity to highly synchronized bursts (**Fig. 2F**), as evidenced by a significant increase in the cross-correlation indexes among these units (**Fig. 2H-I**) and a marked decrease in the time lags (**Fig. 2J-K**), which indicate the temporal differences between the peak firing of these units. Our *in vivo* electrophysiology experiments thus present compelling evidence that BK-D434G MLT neurons fire highly synchronized bursting spikes during the ictal phase of absence seizures.

**Figure 2.**
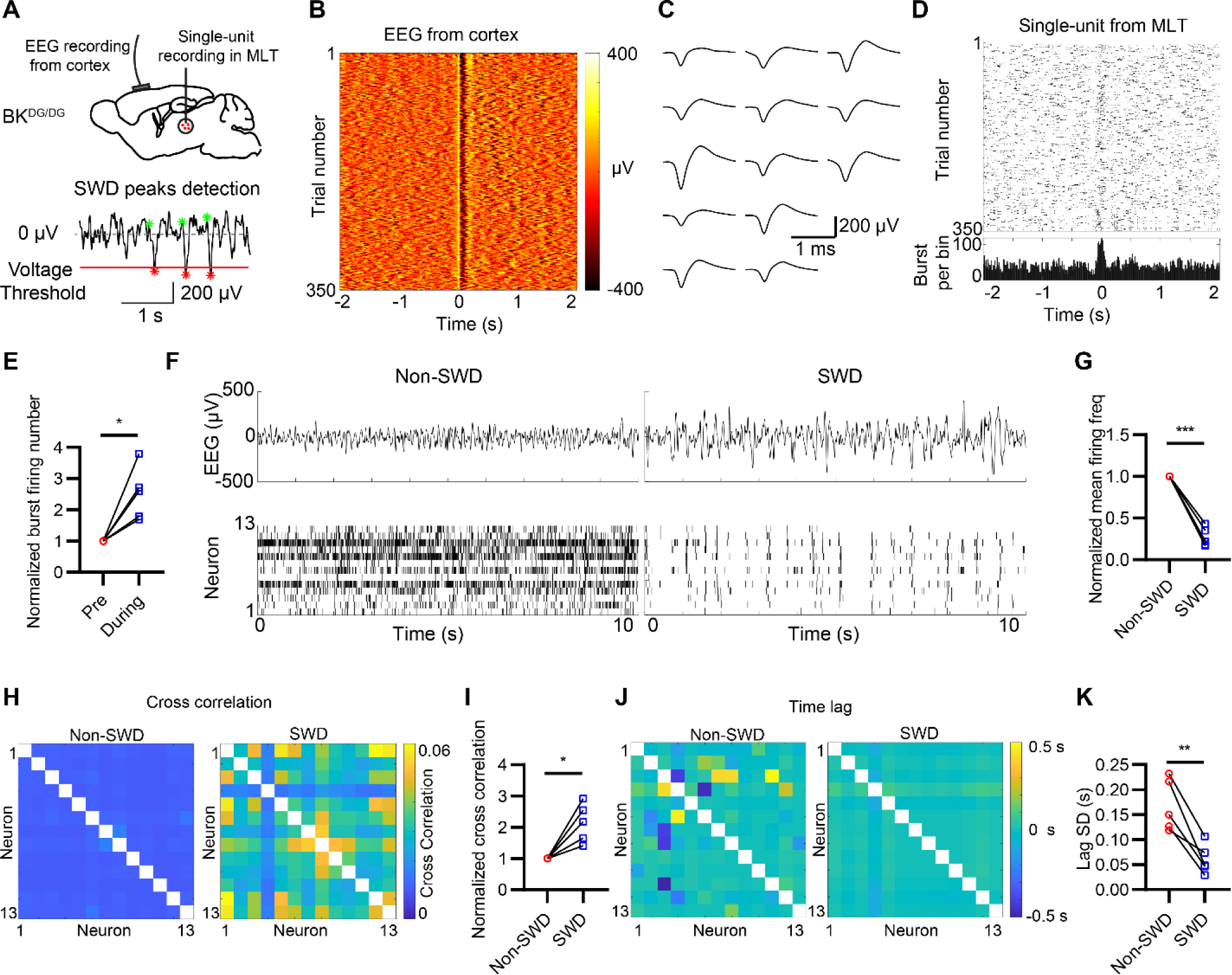
BK-D434G midline thalamus neurons (MLT) exhibit synchronized burst firing during absence seizures *in vivo*. (A) (Upper panel) Schematic of the simultaneous electroencephalogram (EEG) recordings with the single-unit recording in the MLT of freely moving BK^DG/DG^ mice and (Lower panel) the voltage threshold method to detect the onset time of SWD from EEG recording (see Fig. S3 and Methods). (**B**) Representative heatmap of EEG aligned to the peaks of SWDs. (**C**) Averaged spike waveform of individual MLT neurons recorded by single unit electrodes. (**D**) (Top) Representative raster plot of MLT burst firing (> 150 Hz) recorded from the BK^DG/DG^ mice *in vivo*. The single-unit signals were aligned to the onset of SWD. (Bottom) Peri-stimulus time histogram (PTST) of the MLT burst firing. The PSTH shows the summary of all the recorded MLT burst firing during all the 350 SWD peaks. (**E**) Relative change of burst firing number before (pre) or during the SWD peaks, each pair of dots is the mean burst firing number of all the neurons of one mouse. Two-tailed paired Student’s *t*-test, t_4_ = 3.991, *P* = 0.0163. (**F**) Representative raw EEG traces (top) and *in vivo* MLT firing (bottom) during a non-SWD (inter ictal) and a SWD (ictal) period. (**G**) Relative change of mean firing frequency normalized to the non-SWD period. Two-tailed paired Student’s *t*-test, t_4_ = 14.06, *P* = 0.0001. (**H**-**K**) The BK^DG/DG^ MLT neurons exhibit hyper-synchronized burst firing during the ictal period with a higher cross correlation index (**H**-**I**) and a lower time lag among different neurons (**J**-**K**). (**H**) Cross correlation indexes among different neurons during non-SWD and SWD periods. (**I**) Relative cross correlation index normalized to the non-SWD period. n = 5 mice. Two-tailed paired Student’s *t*-test, t_4_ = 4.081, *P* = 0.0151. (**J**) Mean time lag among different neurons during non-SWD and SWD periods. (**K**) Standard deviation (SD) of time lag among different neurons during non-SWD and SWD periods. n = 5 mice. Two-tailed paired Student’s *t*-test, t_4_ = 5.367, *P* = 0.0058.

### BK-D434G MLT neurons exhibit enhanced bursting spikes *ex vivo*

T-CaV channels enable low-threshold spikes (LTS) and subsequent burst firing in thalamic neurons^32^. BK channels can functionally couple to T-CaV channels owing to their spatial close proximity within the calcium channel nanodoamin^33^. Given that BK-D434G has increased calcium sensitivity^15, 34, 35^, the gain-of-function mutation may enhance its functional coupling to T-CaVs and alter the shape and frequency of the T-CaV-mediated rebound burst firing in the MLT neurons. To test this hypothesis, we measured the MLT LTS using whole-cell current clamp (**Fig. 3A**). A −200-pA negative current injection induces LTS-mediated rebound burst firing in BK^WT/WT^ MLT neurons, which is drastically enhanced in BK^DG/DG^ MLT neurons (**Fig. 3B-C**). The burst firing frequency of BK^DG/DG^ MLT neurons is significantly higher than that of BK^WT/WT^ MLT neurons, supported by a notable reduction in the inter-spike interval (ISI) (**Fig. 3B & D**). Overlaying the first LTS spike revealed that BK^DG/DG^ MLT neurons exhibit much faster repolarization as evidenced by the significantly shortened action potential duration (AP90) and augmented after-hyperpolarization amplitude (AHP) (**Fig. 3B, E, F**). As K^+^ efflux through BK channels contributes to fast after-hyperpolarization (fAHP)^36, 37^, our observation of accelerated repolarization and enhanced AHP in the *BK^DG/DG^* neurons suggests that the gain-of-function BK-D434G mutant channels, which have a higher calcium sensitivity^15, 35^, become more active following T-CaV-mediated calcium entry and membrane depolarization during LTS. Owing to its large conductance, potassium efflux through BK-D434G channels rapidly repolarizes the MLT neurons (**Fig. 3E**) and enhances AHP (**Fig. 3F**), which enables rapidly deinactivation of the voltage gated sodium channels (NaVs) and subsequent enhanced rebound bursting^14^. Taken together, our *ex vivo* current clamp characterization shows that BK-D434G promotes LTS-mediated bursting spikes in MLT neurons, which may contribute to the hypersynchronicity of MLT bursting observed in BK^DG/DG^ mice *in vivo* (**Fig. 2**).

**Figure 3.**
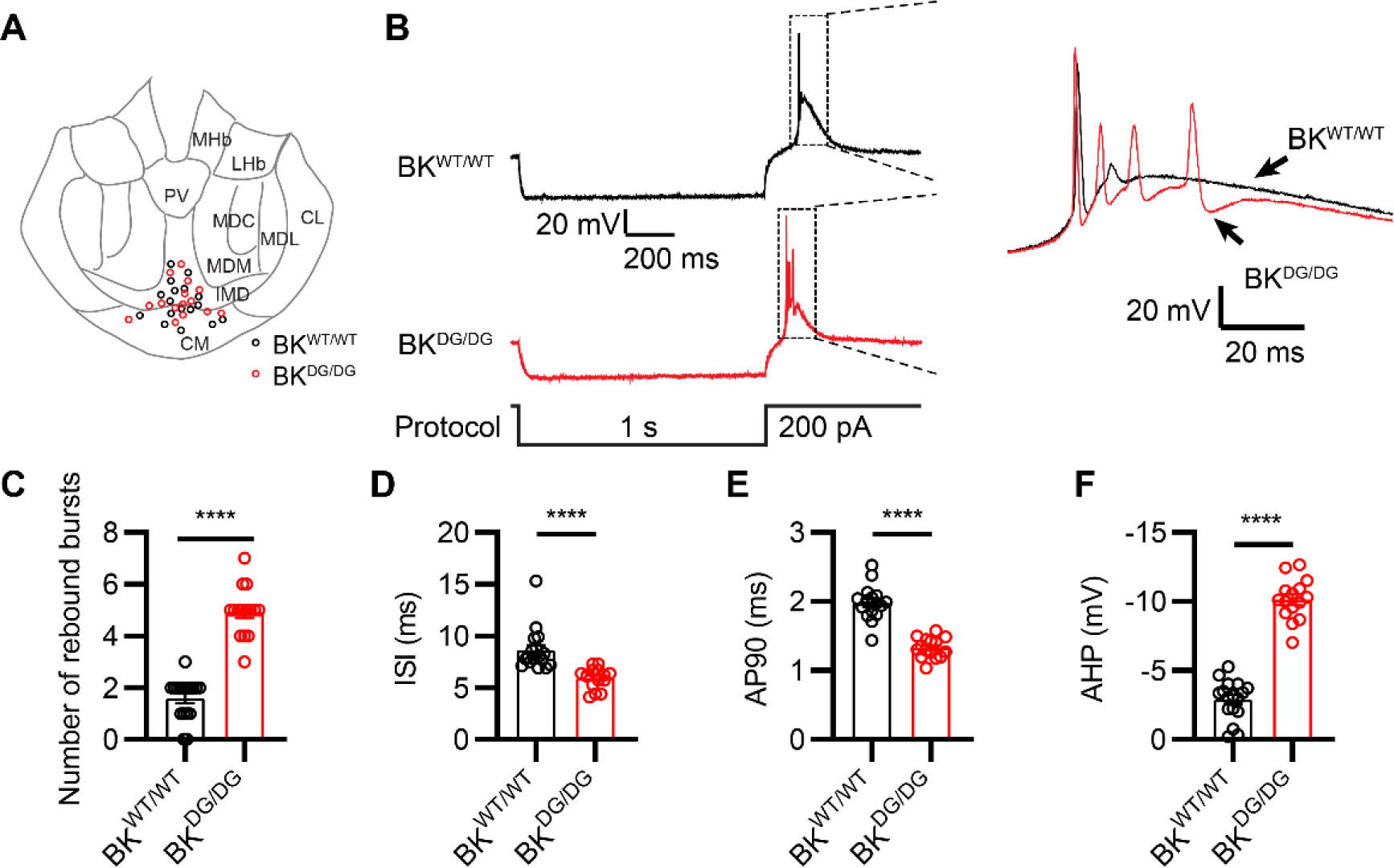
BK-D434G MLT neurons show increased hyperpolarization-induced, low-threshold burst firing *ex vivo*. (**A**) Recording sites across the different subregions of MLT from the BK-D434G mice (BK^WT/WT^ neurons: black circles; BK^DG/DG^ neurons: red circles). (**B**) Representative hyperpolarization-induced rebound burst firing from the MLT neurons (left) and the overlay of the burst firing on an expanded time scale (right). The rebound burst firing was elicited by a −200 pA current injection for 1 s (bottom). (**C**-**F**) Quantification of the rebound burst firing from BK^WT/WT^ and BK^DG/DG^ MLT neurons: number of rebound burst firing (**C**, two-tailed unpaired Student’s *t*-test, t_30_ = 10.77, *P* < 0.0001), inter-spike-interval (**D**, two-tailed unpaired Student’s *t*-test, t_30_ = 4.619, *P* < 0.0001), action potential duration (AP90, **E**, two-tailed unpaired Student’s *t*-test, t_30_ = 8.940, *P* < 0.0001) and after-hyperpolarization (AHP, **F**, two-tailed unpaired Student’s *t*-test, t_30_ = 14.01, *P* < 0.0001). n = 17 neurons from four BK^WT/WT^ mice and n = 15 neurons from three *BK^DG/DG^* mice

### Pharmacological inhibition of MLT bursting suppresses absence seizures

Systemic administration of paxilline (PAX), a specific BK channel blocker, efficiently suppresses frequent absence seizures in BK-D434G mice^14^. To test whether PAX targets the BK channels in the MLT neurons to mitigate absence seizures, we conducted three sets of experiments. First, we employed *in vivo* single-unit recording and monitored MLT neuronal firing before and after systemic PAX administration. We found that PAX effectively abolishes synchronous burst firing in the BK^DG/DG^ MLT (**Fig. S4A**), as evidenced by increased overall firing rate (**Fig. S4B**), decreased cross correlation indexes (**Fig. S4C-D**), and increased time lags (**Fig. S4E-F**) between different units. Our finding suggests that PAX may directly target MLT BK channels and inhibit synchronous burst firing. Next, we measured the direct effect of PAX on LTS-mediated bursting spikes in MLT neurons using brain slice patch clamp recording. PAX significantly reduces LTS-mediated bursting in BK^WT/WT^ and BK^DG/DG^ MLT neurons (**Fig. S5**), indicating that PAX inhibits MLT BK channels and effectively suppresses LTS-mediated bursting. To test the specific effect of PAX on the MLT *in vivo*, we locally infused PAX into the MLT of *BK^DG/DG^* mice through a pre-implanted cannula (**Fig. S2C-D**). We find that, similar to systemic PAX administration^14^, local PAX administration in the MLT also robustly abolishes SWDs and increases vigilance and locomotor activity of the freely behaving mutant mice (**Fig. S6**). Our *ex vivo* and *in vivo* observations with PAX thus support that pharmacological suppression of LTS-mediated bursting in MLT neurons may be a promising approach to treat absence seizures.

### Optogenetic stimulation prevents MLT bursting and suppresses absence seizure

Next, we tested whether MLT burst firing and absence seizures in BK^DG/DG^ mice can be attenuated by optogenetic manipulation, which has higher spatiotemporal resolution than pharmacological approaches. First, we conducted *ex vivo* current clamp recording on ChR2-expressing BK^DG/DG^ MLT neurons (**Fig. 4A**). When the MLT neurons were held at −60 mV at which most of the T-CaVs are inactivated^38^, a 20 Hz light stimulation entrained neuronal firing into tonic mode with high fidelity (**Fig. 4B-C**). No burst firing was elicited. When the membrane was held at −85 mV to deinactivate T-CaVs, the 20 Hz light stimulation first elicited transient burst firing (∼150 Hz) followed by a rapid transition to tonic firing mode, which was entrained at 20 Hz (**Fig. 4B-C**). Our *ex vivo* recording demonstrates that the MLT neurons, similar to other thalamocortical neurons, can switch firing modes between tonic and bursting^2, 5, 11^. Regardless of their membrane potentials, tonic optogenetic stimulation can rapidly entrain BK^DG/DG^ MLT neurons in tonic firing mode and prevents their burst firing.

**Figure 4.**
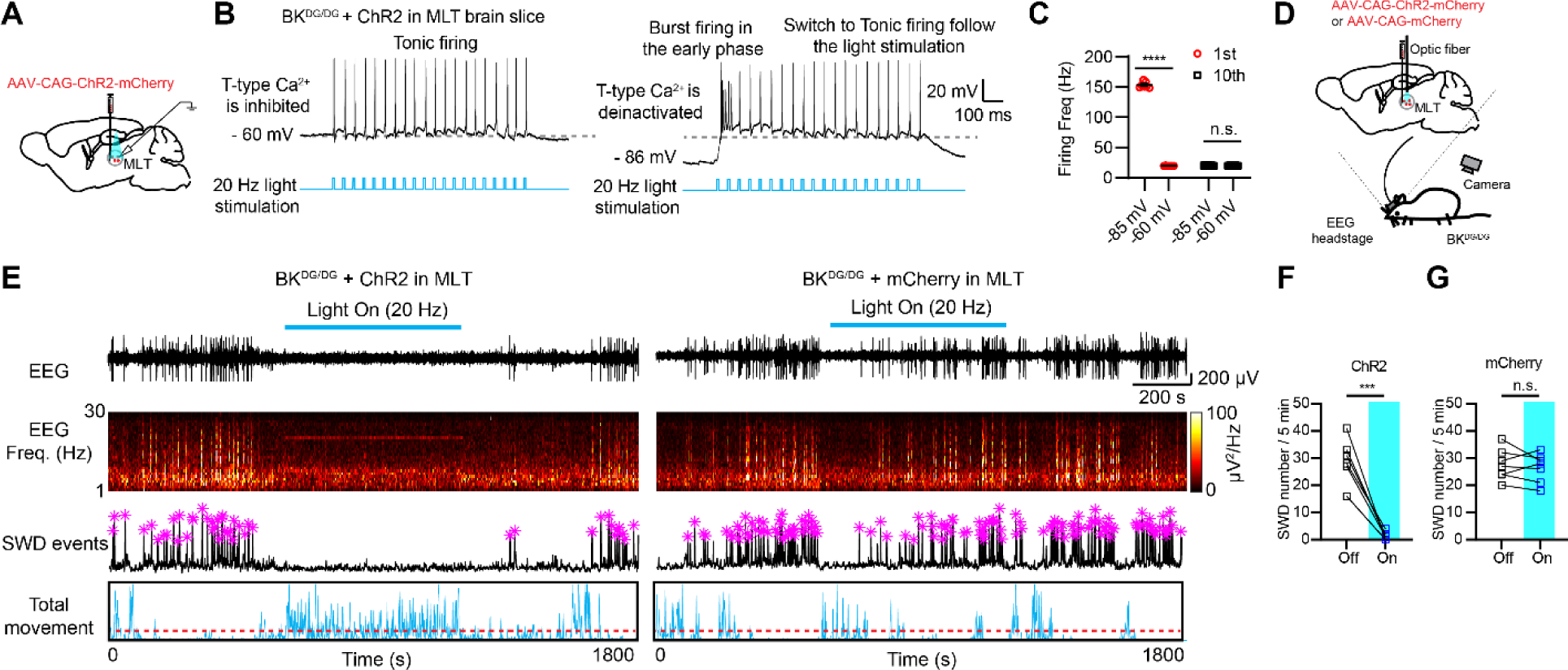
Optogenetic switching of BK-D434G MLT firing mode from bursting to tonic suppresses spontaneous absence epilepsy. (**A)** Schematic of the experimental setup for *ex vivo* MLT patch clamp recording with optogenetic stimulation. **(B**) Representative light-evoked spikes of ChR2-expressing MLT neurons from BK^DG/DG^ mice at holding membrane potential of −60 mV (left) or −86 mV (right). A 5 ms blue light stimulation was delivered at 20 Hz for 1 s (bottom). (**C**) Comparison of the firing frequency induced by the 1^st^ or 10^th^ light stimulation when the membrane potential was held at −60 mV or – 85 mV. Two-way ANOVA, *F*_(1,12)_ = 4731, *P* < 0.0001. n = 7 neurons). (**D**) Schematic of the experimental setup for *in vivo* MLT optogenetic stimulation (see Methods for details). (**E**) Video-EEG recording of BK^DG/DG^ mice in response to a 20 Hz tonic optogenetic stimulation in the MLT infected with ChR2 (left) or mCherry (right). From top to bottom, raw EEG traces, corresponding spectrograms of the EEG traces after fast Fourier transformation (FFT), EEG intensity of EEG with the purple asterisks labeling the SWD peaks, video-based analysis of the total movement that reflects behavioral arrest or vigilance level. (**F, G**) Quantification of the effects of the tonic optogenetic stimulation on the SWD number for BK^DG/DG^ mice expressed ChR2 (**F,** Two-tailed paired Student’s *t*-test, t_6_ = 7.829, *P* = 0.0002) or mCherry in the MLT (**G,** Two-tailed paired Student’s *t*-test, t_6_ = 0.7233, *P* = 0.4967). n = 7 mice per group.

Subsequently, we examined whether the optogenetic modulation of MLT firing mode can suppress absence seizures in freely moving BK^DG/DG^ mice (**Fig. 4D**). Viral expression ChR2 or control mCherry in the MLT of BK^DG/DG^ mice (**Fig. S2E-F**) does not alter their frequent SWDs and behavioral arrest as evidenced by the lack of locomotor activity (**Fig. 4E-G**). A single optogenetic stimulation (3 seconds, 20 Hz, 5 ms pulse width) or continuous light stimulation (10 minutes, 20 Hz, 5 ms pulse width) of the ChR2-expressing MLT but not the mCherry-expressing MLT immediately suppresses SWD generation and rescues the animals from seizure-induced behaviorally arrested state, dramatically promoting locomotor activity and vigilance (**Fig. 4E-G, Video S1**). Our *ex vitro* and *in vivo* optogenetic experiments thus collectively show that entraining MLT neurons in tonic firing mode prevents their burst firing and effectively suppresses absence seizures in BK^DG/DG^ mice. Immediate rescue from the behaviorally arrested state and increase of locomotor activity of BK^DG/DG^ mice upon light stimulation further demonstrate the important role of the MLT in regulating vigilance and absence seizures.

### Vigilance enhancement suppresses MLT bursting and prevents absence seizure

Clinical evidence has established the inverse relationship between vigilance level and the incidence of absence seizures, but with an unclear mechanism^29–31^. To dissect the role of the MLT in mediating the inverse correlation and explore an alternative therapeutic strategy to mitigate absence epilepsy, we examined the impacts of amphetamine, a commonly used stimulant^39, 40^, on the vigilance, absence seizures, and MLT firing of BK^DG/DG^ mice. Systemic administration of 3 mg/kg D-amphetamine sulfate dramatically suppresses absence seizures and increases the vigilance of the animals as evidenced by their enhanced locomotor activity compared to saline control (**Fig. 5A-B**). Further, no SWD was detected while the mice constantly moved due to the stimulating effect of amphetamine. Our single-unit *in vivo* recording from the MLT further demonstrates that amphetamine administration increases overall firing rate (**Fig. 5C-D**) and desynchronizes the MLT neuronal firing as evidenced by the decreased cross correlation indexes (**Fig. 5E-F**) and increased time lags (**Fig. 5G-H**) between different units. The desynchronization of MLT burst firing recapitulates the changes of *in vivo* multiunit signals observed with systemic administration of PAX (**Fig. S4**). Interestingly, the anti-absence seizure effect of D-amphetamine lasts much longer than PAX. Consistent with our recent observation that D-amphetamine has no direct effect on BK channel^41^, we find that it does not directly change the LTS burst firing in BK^DG/DG^ MLT neurons (**Fig. S7**). Our amphetamine experiments thus suggest that the stimulant may work on the brain regions that innervate the cortical-TRN-thalamic system including the MLT, thereby indirectly suppressing the bursting of thalamocortical neurons and preventing absence seizures. Our amphetamine administration (**Fig. 5**) and the treadmill perturbation (**Fig. 1D-J**) together indicate that the MLT plays a key role in regulating vigilance and absence seizures. Vigilance enhancement, likely through an indirect, circuit-based mechanism, suppresses the synchronized burst firing of the BK^DG/DG^ MLT neurons and prevents absence seizures.

**Figure 5.**
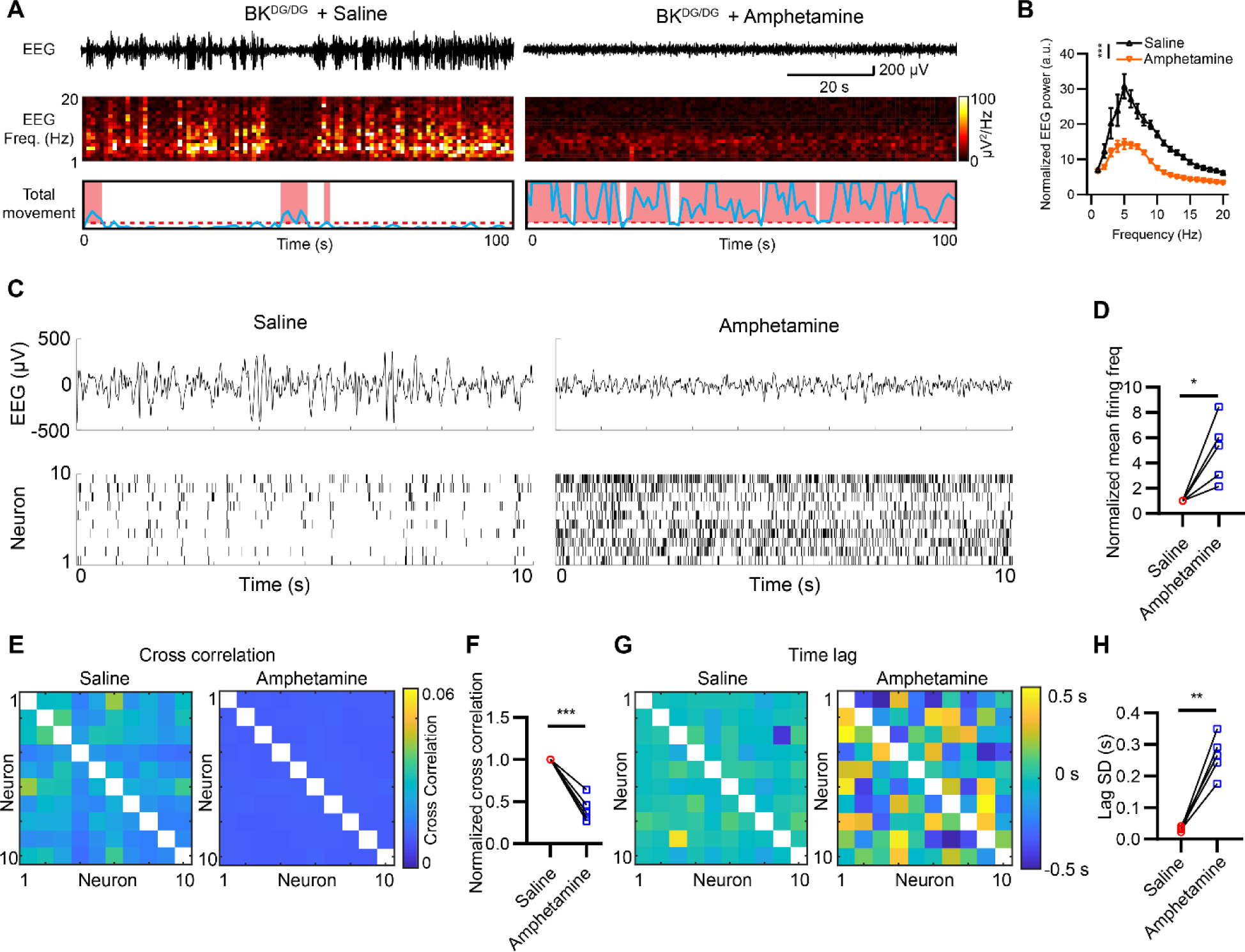
Amphetamine suppresses absence seizures and desynchronizes MLT burst firing *in vivo*. (**A**) Representative EEG traces (top), corresponding spectrograms (middle), and total movement (bottom) from a BK^DG/DG^ mouse before and after 3 mg/kg D-amphetamine sulfate administration. (**B**) Summary of power spectral density of EEG recorded from *BK^DG/DG^* mice before and after amphetamine administration. Two-way ANOVA, *F*_(1,8)_ = 31.07, *P* = 0.0005. n = 5 mice. (**C**) Representative raw EEG traces (top) and single-unit signals (bottom) from the BK^DG/DG^ MLT neurons of the same animal after saline (left) and amphetamine (right) administration. (**D**) Relative change of mean MLT neuronal firing frequency after amphetamine administration. The firing frequency was normalized to the SWD period of the saline group. Two-tailed paired Student’s *t*-test, t_4_ = 3.588, *P* = 0.0230. (**E**-**H**) Hyper-synchronized burst firing of BK^DG/DG^ MLT neurons diminishes after amphetamine administration. (**E**) Cross correlation indexes among different neurons after saline and amphetamine administration. (**F**) Relative cross correlation indexes normalized to the saline group. Two-tailed paired Student’s *t*-test, t_4_ = 8.971, *P* = 0.0009. (**G**) Mean time lag among different neurons after saline and amphetamine administration. (**H**) Standard deviation of time lag among different neurons after saline and amphetamine administration. Two-tailed paired Student’s *t*-test, t_4_ = 7.957, *P* = 0.0014. n = 5 mice. In all plots and statistical tests, summary graphs show mean ± s.e.m.

### Deep brain stimulation of the MLT suppresses absence seizures in BK-D434G mice

Deep brain stimulation (DBS) offers spatiotemporal precision and versatility to modulate neuronal activity. It has been successfully applied to treat various neurological disorders, including epilepsy, essential tremor, and Parkinson’s disease^42^. Given the spatial confinement of the MLT, we reasoned DBS modulation of MLT firing mode could be a feasible strategy to treat absence epilepsy. To test this hypothesis, we implanted a stimulation electrode array into the BK^DG/DG^ MLT (**Fig. S2G-H**). To reliably control absence seizures, we implemented a closed-loop DBS system, which automatically detects SWDs *in situ* and immediately delivers a 20 Hz, 50 µA, 1 ms pulse width DBS command for 5 seconds to the MLT in real-time (**Fig. 6A**, see Methods). Our closed-loop DBS system accurately detects the SWD peaks and triggers transistor–transistor logic (TTL) signals to deliver DBS to the MLT (’on’, **Fig. 6E**). With DBS, the typical 3∼8 Hz absence seizure bands were suppressed, and the vigilance level of the same animal immediately increased, accompanied by enhanced locomotor activity and a nearly complete suppression of SWDs (**Fig. 6B-E**, **Video S2**). On the other hand, in the control group, where DBS was not triggered (‘off’, **Fig. 6D**), the BK^DG/DG^ mice developed spontaneous absence seizures with typical 3-8 Hz absence seizure EEG frequency band and frequent behavioral arrests (**Fig. 6B-D**). We did not observe detectable adverse effects such as abnormal postures during closed-loop DBS (**Video S2**). Our closed-loop DBS experiments further support that the MLT plays a key role in regulating absence seizure genesis and vigilance level. Electrical modulation of the MLT through closed-loop DBS is a promising strategy to treat absence epilepsy.

**Figure 6.**
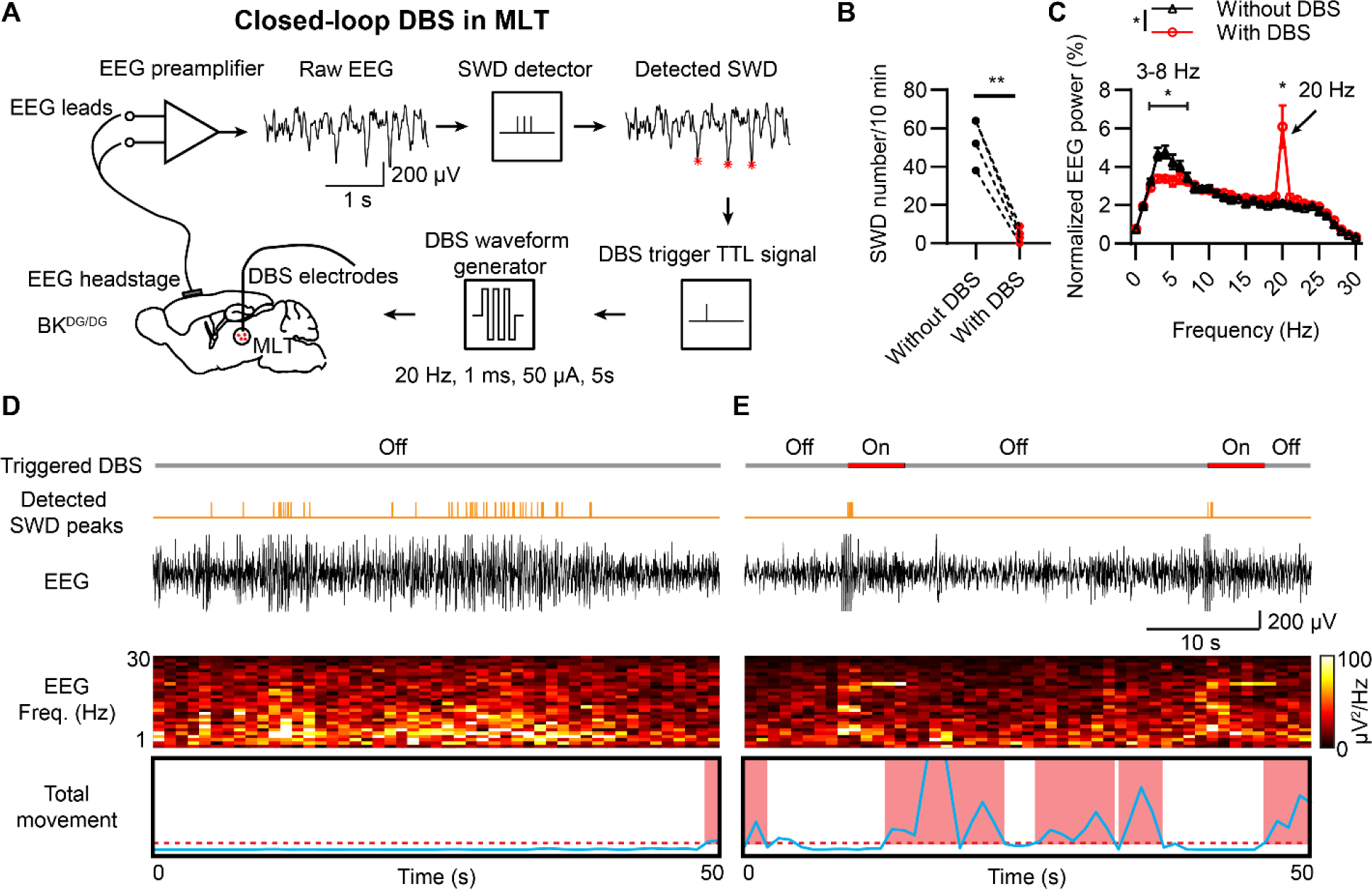
Closed-loop deep brain stimulation (DBS) of the midline thalamus attenuates spontaneous absence seizures in BK-D434G mice. (**A**) Diagram of a closed-loop DBS system. SWDs are detected using the algorithm to calculate the SWD indexes from amplified EEG signals in real time (see Methods). Once an SWD event is detected, a 5-s electrical stimulation (biphasic rectangular pulses with a pulse width of 1 ms, a current power of 50 µA, a frequency of 20 Hz) is generated and immediately delivered to the MLT via a DBS electrode array. (**B-C**) Effects of the closed-loop DBS on the number of SWDs (B, Two-tailed paired Student’s t-test, t_3_ = 10.99, P = 0.0016) and EEG power (C, Two-way ANOVA, interaction, F = 9.118, P = 0.0172) of BK^DG/DG^ mice over a 10-minute period. Each group consists of 4 mice. Notably, the DBS group shows a significantly reduction in the SWD band from 3-8 Hz (Two-way ANOVA, F_(1, 3)_ = 22.44, P = 0.0178), whereas the power at stimulation frequency of 20 Hz increases during DBS (arrows, Two-tailed paired Student’s t-test, t_3_ = 3.758, P = 0.0329). (**D-E**) Representative video-EEG results of the same BK^DG/DG^ mouse with (**E**) and without (**D**) the closed-loop DBS in a 50-second period. From top to bottom: DBS protocol, detected SWD peaks (orange spikes), raw EEG traces, EEG power, and total movement based on video analysis.

## Discussion

In this study, using a new absence epilepsy mouse model with direct human relevance, we gained critical insights into the role of the MLT nuclei in absence seizure pathogenesis, the underlying ionic basis of enhanced MLT bursting, and potential strategies to suppress MLT neuronal bursting to treat absence epilepsy (**Fig. 7**).

**Figure 7.**
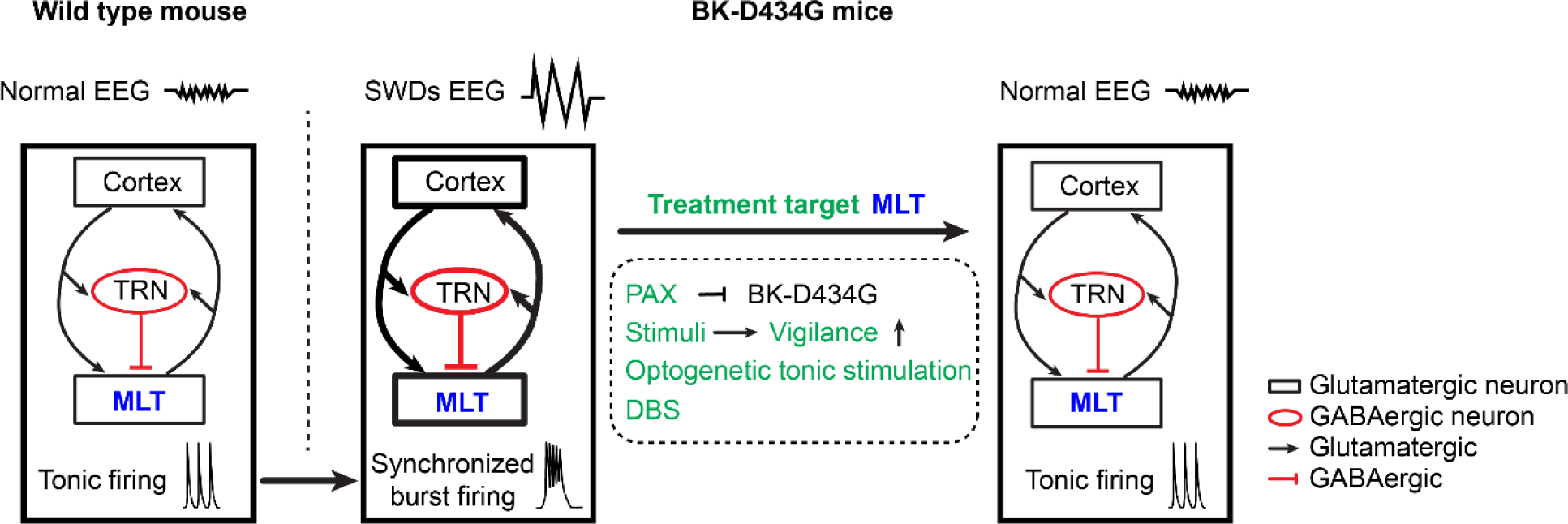
A mechanistic diagram depicts the role of midline thalamus (MLT) bursting in absence seizure pathogenesis and outlines new anti-absence interventions by targeting MLT bursting. BK-D434G channelopathy disrupts the balance within the cortico-thalamic reticular nucleus (TRN)-MLT circuitry, promoting hypersynchronized bursting in the MLT neurons and absence seizures. Suppressing MLT hypersynchronization by BK channel inhibition (paxilline or PAX), vigilance enhancement by physical perturbation or amphetamine administration, optogenetic stimulation or deep brain stimulation (DBS) of the MLT, effectively prevents SWD generation and absence seizures.

Our current study fills a critical knowledge gap with regard to the MLT’s role in absence seizure pathogenesis. The MLT is a higher-order thalamic region that projects broadly to large cortical areas and regulates vigilance and arousal^25, 26^. The seminal work done by Jasper and colleagues in the 1940s and 1950s first hinted at the involvement of the MLT in absence epilepsy^27, 43^. The authors showed that low-frequency electrical stimulation of the MLT in feline models can induce absence seizure-like phenotypes. On the other hand, high-frequency electrical stimulations led to desynchronized cortical activity, potentially promoting arousal and viligance^27^. Despite these pioneering findings, the exact role of the MLT in absence seizure pathogenesis has not been established in other animal models using contemporary technologies and tools. Utilizing the BK-D434G absence seizure mouse model, we discovered that the MLT neurons show hypersynchronized bursting during the ictal phase. Specific modulations of MLT bursting, including local infusion of BK channel antagonist PAX, MLT-specific optogenetic stimulation and closed-loop DBS, effectively suppress SWDs, increase vigilance, and eliminate frequent spontaneous absence seizures (**Fig. 7**). Our findings thus explicitly establish that the MLT is a critical thalamic region, whose hypersynchronized burst firing contributes to absence seizure propagation and maintenance.

Establishing the MLT as a key thalamic region for absence seizure pathogenesis helps understand the inverse relationship between absence seizures and vigilance levels, a hallmark of absence epilepsy^30, 44^. While absence seizures occur more frequently during drowsy or passive awake state^29^, increased vigilance induced by playing with novel toys or participating challenging tasks effectively prevents absence seizures^30^. The MLT is known to be an important regulatory center for vigilance and arousal^25–28^. We found that there is increased c-Fos expression in the MLT that corresponds with the absence seizures in BK-D434G mice (**Fig. 1**). The pattern of this c-Fos expression in the BK-D434G MLT mirrors what was observed in another rodent model for absence seizures, the WAG/Rij rat, in which the seizures were triggered by hypoxia-induced hyperventilation^45^. These findings suggest that the role of the MLT in absence seizures is not limited to the context of the BK-D434G mutation. Instead, it could represent a potential general mechanism for absence seizures with different etiologies. Our specific modulations of MLT bursting in freely behaving BK-D434G mice during seizure attacks unexceptionally led to immediate cessation of SWDs and increased of vigilance as evidenced by terminating the behavioral arrested state and concomitant increase of locomotor activity (**Figs. 4-6**). In addition, enhancing vigilance by forced movement on the treadmill (**Fig. 1E-G**) or amphetamine (**Fig. 5**) robustly eliminates absence seizures with direct interference with MLT firing as evidenced by the decreasing the c-Fos expression (**Fig. 1H-J**) or desynchronized firing (**Fig. 5C-H**). All these findings support the important role of MLT bursting in regulating vigilance/arousal levels and absence seizures (**Fig. 7**).

Our *ex vivo* brain slice electrophysiology also advances understanding of the ionic basis of neuronal burst firing in the MLT. It is well known that T-CaV channels are responsible for the generation of LTS, which subsequently trigger bursting in thalamic neurons^2, 11^. We observed that BK-D434G MLT neurons have markedly enhanced LTS-mediated burst firing with accelerated repolarization, narrowing of action potential duration, enhanced AHP, and shortened inter-spike-interval (**Fig. 3B-F**). Inhibiting BK channels with PAX significantly reduces BK-D434G MLT burst firing (**Fig. S5**). This is consistent with the effectiveness of PAX, either administered locally to the MLT (**Fig. S6**) or systemically^14^, in suppressing absence seizures. Our study thus shows that BK channels play an important role in facilitating LTS-mediated burst firing. It is likely that T-CaV-mediated calcium entry and membrane depolarization during LTS more efficiently activate the gain-of-function mutant BK channels. This leads to a more robust BK-mediated K^+^ efflux, stronger membrane repolarization, likely rapid de-inactivation of the voltage-gated sodium channels, and subsequent increase of burst firing frequency. Future studies are required to further test this model.

Our *in vivo* single-unit recording of MLT neurons shines light on understanding the contributions of these understudied thalamocortical neurons to absence seizure pathogenesis. In addition to the hypersynchronized burst firing in BK-D434G MLT neurons, we also detected a decrease in the overall firing rate of these neurons during the ictal phase (**Fig. 2F-G**). This reduction in neuronal activity during the ictal phase is consistent with findings from another rodent model of absence seizures^46^. Interestingly, there is always a transient, pre-ictal silent phase followed by dramatic increase of bursting at the onset of SWDs (**Fig. 2D-E**). The reduction of overall firing rate and the sudden silencing of the MLT neurons likely reflect the concomitant reduction of excitatory inputs from the other brain regions that project to the MLT and enhanced inhibitory inputs from TRN neurons (**Fig. 7**). Both effects may potentially help hyperpolarize the MLT membrane, facilitating de-inactivation of T-CaVs and promoting LTS and subsequent bursting.

On the contrary, systemic administration of amphetamine (**Fig. 5**) or PAX (**Fig. S4**) to BK-D434G mice effectively increases MLT overall firing rate and desynchronizes the MLT firing accompanied by diminished SWDs and suppression of absence seizures. Our *ex vivo* and *in vivo* electrophysiology experiments thus collectively demonstrate that the intrinsic properties of MLT neurons that control burst firing, including T-CaVs and BK channels, and the extrinsic factors that affect MLT membrane potential to enable T-CaV-mediated LTS and bursting, including direct and indirect synaptic inputs to MLT neurons, are critical in determining MLT bursting and absence seizure pathogenesis (**Fig. 7**).

Based on this model, we tested several complementary approaches to suppress MLT bursting, which have translational potentials to treat absence epilepsy with different etiologies, including drug resistant absence epilepsy. First, pharmacological inhibition of BK channels in the MLT neurons is an effective and direct approach to suppress MLT bursting and prevent seizures (**Fig. S5-6**). However, the anti-absence effect of paxilline, either systemically^14^ or locally administered (**Fig. S6E**), does not persist *in vivo*. Discovering BK antagonists with better pharmacokinetics is needed to further develop this new anti-absence strategy.

In addition, enhancing vigilance is known to be an indirect yet effective approach to suppress MLT bursting and prevent absence seizures (**Fig. 5**). In fact, amphetamine, a widely used stimulant medication, was used to manage absence seizures^47, 48^ before ESM was developed in the late 1950s^49^. Amphetamine was then replaced by ESM in treating absence epilepsy due to its addictive nature. We showed that amphetamine does not directly inhibit BK channels^41^ and it has no direct effect on MLT bursting (**Fig. S7**). Therefore, the stimulant likely exerts its anti-absence effect through an indirect mechanism to influence MLT bursting. Future studies are needed to dissect the underlying mechanism and to test the effectiveness of less-addictive stimulants on preventing absence seizures. Interestingly, we found that physical perturbation such as low-speed treadmill also effectively reduces MLT hyperactivity, increases vigilance level, and prevents absence seizures (**Fig. 1E-G**). We hypothesize that physical perturbation may strengthen sensory inputs to the MLT, a vigilance and arousal center. This can depolarize MLT neurons thereby preventing hypersynchronized LTS-mediated bursting. Although the exact mechanisms underlying these vigilance enhancement approaches demand further investigations, our alternative approaches may inspire new antiabsence therapeutics with less adverse effects than pharmacological approaches.

Finally, precise modulation of MLT bursting with optogenetics and DBS is another effective approach to prevent absence seizures (**Figs. 4 and 6**). By switching the firing mode of MLT neurons from bursting to tonic, optogenetic stimulation immediately increases vigilance and abolishes absence seizures. While optogenetic techniques have been widely employed in basic research, and Adeno-Associated Virus (AAV) treatments have received approval for early-phase human trials, the clinical applications of optogenetics are still in their nascent stages. In contrast, DBS has been successfully applied to treat various neurological disorders, including dystonia, epilepsy, essential tremor, and Parkinson’s disease^42^. This success can be attributed to DBS’s ability to modulate neuronal firing, brain oscillations, and synaptic plasticity. We showed that closed-loop DBS of the MLT can effectively inhibit absence seizures and rescue behavioral arrest in BK-D434G mice (**Fig. 6**). We believe that, after further optimization of DBS parameters, closed-loop DBS on the MLT may be a promising anti-absence therapeutic strategy to treat absence epilepsy, specifically pharmaco-resistant absence epilepsy.

In summary, our characterizations of BK-D434G mouse model demonstrate that MLT bursting plays a crucial role in absence seizure pathogenesis. The molecular and circuit mechanisms of MLT bursting identified in this study and the mechanism-derived anti-absence approaches advance our mechanistic understanding of absence seizure pathogenesis and will inspire future development of therapeutics to treat pharmaco-resistant absence epilepsy.

## Methods

### Mice

The generation of BK-D434G mutation mice was previously described^14^. PCR genotyping was performed using tail DNA extraction. The genotyping was performed by using the genotyping primers *pKCNMA1*_genotyping F1 (5’-GTGCCTAGAGGTGGCTGGGAATTAG-3’) and *pKCNMA1*_genotyping R1 (5’-CCTCTCCTACGGTGGTAAAGTATCC-3’). The length of wildtype band is 342 base pairs (bp), and the mutated band is 455 bp. The heterozygous BK-D434G (*BK^DG/WT^*) males were bred with the *BK^DG/WT^*females. All mice were housed in a 12-h light-dark cycle (light on at 07:00 am) with *ad libitum* food and water. The usage and handling of the mice are conducted with strict compliance with protocols approved by the Institutional Animal Care and Use Committee at Duke University and National Institutes of Health guidelines.

### Immunostaining

c-Fos was used as a marker to report absence seizure-induced neuronal activities, both firing rates and firing patterns^17–19^. To measure spontaneous seizure-induced c-Fos expression, BK-D434G mice were allowed to habituated for 1 hour. As previously reported^14^, the animals develop frequent absence seizures (>10 SWD per 5 min) accompanied by behavioral arrest after habituation. Given that c-Fos protein takes approximately 2.5 hours to reach maximum expression post certain behaviors^20^, the animals were euthanized 2.5 hours after the 1-hour habitation period. WT littermates, used as controls, were subjected to the same conditions. For the forced movement on treadmill experiments, BK-D434G mice were placed on a treadmill, either running or stationary. The animals were euthanized 2.5 hours after the treadmill section. The brain tissues were fixed using transcardial perfusion of cold PBS followed by 4% paraformaldehyde. We collected 50 μm coronal sections containing the MLT using a cryostat (Leica CM1900). After rinsing the sections three times with PBS for 10 min each, we blocked them with 5% goat serum and 0.3% Triton X-100 for 2 hours at room temperature. The sections were then incubated overnight at 4°C with the primary c-Fos antibody (1:1000, Rabbit, Cell Signaling #2250). After 3 rinses with PBS for 10 min, secondary antibodies (1:1000, conjugated with Goat anti-rabbit Alexa 488, Thermo Fisher Scientific, A-11008) were incubated for 2 hours at room temperature. Then, the sections were washed 3 times with PBS for 10 min each and stained with DAPI (1:10000 of 5 mg/mL, Sigma-Aldrich) before mounted. Images were acquired using a Zeiss 780 inverted confocal microscope. Representative images were acquired from at least three biological repeats. We analyzed the number of c-Fos-labeled cells using a custom-written MATLAB program (https://github.com/superdongping/c_fos_count).

Following similar immunostaining procedure above, histological analysis was done for all single-unit recording, optogenetic stimulation, cannula drug delivery, and DBS experiments to confirm the implantation and/or viral injection sites. After imaging, all the detected sites were recorded and registered in a brain atlas.

### Video-EEG recording

Video EEG recording were performed similarly to our previous publication^14^. Briefly, 2 to 6 month-old mice were anesthetized with 1∼2% isoflurane and mounted on a stereotaxic device (Kopf Instruments). An EEG headstage (#8201, Pinnacle Technology Inc., Lawrence, KS, USA) was mounted to the skull by two screws with leading stainless wires on the parietal cortex (from the bregma: 1.50 mm posterior, 2.50 mm lateral) and cerebellar cortex (from the lambda: 1.50 mm posterior, 1.50 mm lateral). The headstage was further secured on the skull by an additional thin layer of super glue (Loctite Inc.) and dental cement. EEG recording started after a minimum of 7-day recovery period. To record the EEG signals from the cortex, the EEG headstage was connected to a preamplifier with 100X gain and 1 Hz high pass filter (#8200-SE, Pinnacle Technology Inc.). The EEG signal was acquired by an analog to digital converter board (PCI 6221; National Instruments, Austin, TX, USA) connecting to a computer. A webcam (Logitech C920 HD Pro) was set on top of the recording chamber to monitor the animal behavior at acquisition rate 24 frames per second, simultaneously.

### Single-unit recording

For single-unit recording in the MLT neurons, the implantation surgery was performed as previously described^50^. Briefly, a craniotomy was performed above the MLT (coordinates relative to bregma: 1.34 mm posterior, 0.00 mm lateral, −3.50 mm in depth). Then, a drivable electrode which contains 16 recording arrays bundled in a cannula (Innovative Neurophysiology, Inc., Durham, NC, USA) was slowly lowered to 0.3 mm above the MLT with −3.20 mm in depth. The single-unit multi electrodes array was grounded with a leading wire with a screw affixed to the skull (from the bregma: 2.00 mm posterior, 3.00 mm lateral). The single-unit electrode was fixed on the skull by two additional screws followed by super glue and dental cement. After a 7-day recovery, the tips of the single-unit drivable electrode were advanced 0.3 mm deeper, and the headstage was connected to a preamplifier CerePlex mu headstage (Blackrock Neurotech, Salt Lake, Utah). The single-unit recording from the MLT neurons were recorded by the Blackrock Cerebus data acquisition system (Blackrock Neurotech, Salt Lake, Utah), including the neuronal spike waveform, timestamp and analog signal from the EEG headstage. The single-unit recording data were processed by the Offline Sorter x64 V3 (Plexon Inc., Dallas, TX, USA) to sort the single unit through an automatic principal component analysis based on their waveform. The classified single unit signals were further confirmed manually to exclude the artificial or electrical noise based on their waveforms. The sorted single units and EEG analog data were exported to the NeuroExplorer (Nex Technologies. Colorado Springs, CO, 80906, USA) for further analysis.

### SWD timestamp detection

A custom-written MATLAB (MathWorks, Natick, MA, USA) code (https://github.com/superdongping/SWD_detection) was used to detect the timestamps of the SWD peaks from the EEG traces (**Fig. S3**), which were subsequently used to align the SWD peaks with single-unit recording from MLT neurons. The SWD peaks were detected according to the following procedure. As the SWDs of the BK-D434G mice are characterized by 3-8 Hz large amplitude electrical oscillations^14^, a negative voltage threshold was empirically set to −300 μV (**Fig. S3A**) to screen the timestamps of all potential SWD peaks (**Fig. S3B**). The weighted SWD_index_ was then calculated by multiplying the average voltage amplitude (A_n_) of three consecutive potential SWD events (n, n+1 and n+2) with the average frequency (the reciprocal of the difference of inter-event time, T_n_) using Equation 1 (**Fig. S3B-C**).

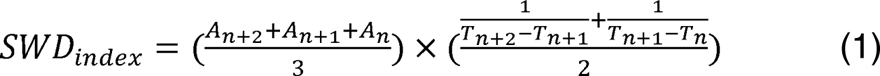

To minimize false positive SWD detection from large-amplitude EEG signals, only the EEG signals with top 75% of SWD_index_ values (SWD_index_ threshold) were included from further analysis (**Fig. S3D**). Red asterisks were used to demarcate the timestamps of the negative peaks that were beyond the SWD_index_ threshold. The timestamps were then overlaid with the raw EEG trace, and the negative SWDs peaks were labeled in the raw EEG trace by the red asterisks (**Fig. S3E**). To find the timestamps of the positive SWD peaks, we first smoothed the raw EEG trace by using the MATLAB build-in function *smooth* (**Fig. S3F**). The SWD positive peaks (green asterisks) were labeled at the maximum positive voltage within the 300 ms time window prior to a negative maximum (red asterisks, **Fig. S3G**). The EEG heatmap was then plotted aligned with the timestamps from the positive SWD peaks.

### Burst firing analysis

To analyze the burst firing activities from the single-unit recorded MLT neurons, a custom MATLAB code was used to calculate the firing frequency of the MLT neurons, and the neuronal firing frequency > 150 Hz was defined as burst firing^51^. A raster plot of MLT burst firing was plotted aligned with the timestamps from the positive SWD peaks. Then a peri-stimulus time histogram (PTST) of the MLT burst firing was used to summarize all the recorded MLT burst firing plot aligned with the timestamps from the positive SWD peaks (https://github.com/superdongping/SWD_detection). The relative change of the burst firing was calculated before or during the SWD peaks. For each mouse, all the burst firing numbers were normalized and averaged together to create population average plots.

### Cross-Correlation analysis

To evaluate the neurons firing synchronicity in the MLT during ictal or inter-ictal period, a custom MATLAB code was written in order to perform cross-correlation analyses among different MLT neurons by utilizing the in-built function *xcorr*. Latencies were obtained by determining the lag of the maximum value of the cross-correlation for positively correlated neurons and the lag to the minimum value for anti-correlated neurons. For each animal, the cross-correlation values compared during ictal or interictal period were summarized and normalized, and all datasets from all animals were averaged together to create population average plots.

### Optogenetic stimulation *in vivo*

For the optogenetic manipulation experiments, surgery was done as described previously^52^. Briefly, 0.3 μl of AAV2/9-CAG-ChR2-mCherry (3 × 10^12^ genomic copies per mL, Addgene, 100054) or AAV2/9-CAG-mCherry (2.85 × 10^12^ genomic copies per mL, Addgene, 108685) was injected into the MLT (from the bregma: 1.34 mm posterior, 0.00 mm lateral, −3.50 mm in depth) of the mice. An optical fiber (200 μm OD, 0.22 NA, Inper Inc., Hangzhou, China) was unilaterally implanted in the MLT (0.2 mm above the viral injection coordinates). An EEG headstage was subsequently implanted on the skull following the same procedure as described above. After the virus was expressed for 3 weeks, the implanted optical fiber was connected to a laser stimulator (Inper Inc., Hangzhou, China) and the laser power at the tip of the fiber was adjusted to ∼3 mW for the blue laser (473 nm). For optogenetic stimulation of the MLT, a 10 min baseline video-EEG was recorded after habituation. Then, a 20 Hz, 5-ms blue light stimulation was delivered to the MLT via the optical fiber for 10 min. The laser power at the tip of the fiber was adjusted to ∼3 mW for the blue laser (473 nm). The mice were monitored for another 10 min after turning off the optical stimulation.

### Pharmacological treatment

Similar to previous descriptions, the mice were allowed to habituate in the recording chamber for 1 hour, which facilitated the development of frequent absence seizures^14^. Subsequently, to evaluate the effect of drug treatment on single-unit firing in the MLT, a single dose of either paxilline (0.35 mg/kg, i.p.), D-amphetamine sulfate (3 mg/kg, i.p.), or a saline control was administered to the mice after recording for 1 hour as a baseline. To test if locally drug infusion can treat absence seizure, a guide cannula (RWD Life Science Inc., Shenzhen, China) was unilaterally implanted into the MLT (from the bregma: 1.34 mm posterior, 0.00 mm lateral, −3.50 mm in depth). The EEG headstage was subsequently implanted additionally on the skull following the same procedure as described above. After 1 week recovery from the surgery, 1 µL, 1 µM PAX, or saline was microinjected into the MLT via an injector (RWD Life Science Inc., Shenzhen, China).

### Locomotor stimulation by treadmill

Each mouse was connected to the pre-amplifier with an EEG headstage before treadmill test. The animals were allowed to freely explore in the treadmill chamber (Treadmill Simplex II, Columbus Instruments, Columbus, OH, USA) for 10 min. Then, the treadmill was turned on and its speed was slowly increased to 2 m/min. After a 10-minute training section, the mice were kept in the treadmill chamber with the treadmill off for 1 hour as baseline. For *BK^DG/DG^*mice, the EEG signals were continuously recorded for 50 min with treadmill on at the speed of 2 m/min followed by another 60 min EEG recording with treadmill off. The animals were simultaneously videotaped at 24 frames per second by a video camera (Logitech C920 HD Pro Webcam), which was placed on the top of the treadmill chamber.

### Closed-loop DBS to treat absence seizure

DBS electrode array (MicroProbes, 2×2 platinum MEA array, spacing 250 µm, impedance 10 kΩ) was stereotaxically implanted into the MLT (from bregma: AP –1.58 mm, ML 0.0 mm, DV –3.50 mm). A headstage (#8201, Pinnacle Technology) was mounted onto the skull to record EEG signals. After the mice recovered from the surgery (> 1 week), they were connected to our customized closed-loop DBS system^53,54^. Once this system detects the onset of SWDs in real-time, it instantaneously responds and delivers a 5-s, 20 Hz, 50 µA, 1 ms pulse width, biphasic DBS to the targeted MLT. Specifically, the EEG signal was routed to an Intan recording system (RHD recording controller, Intan Technologies, sampling rate 10kHz). Custom-written code (Github: superdongping/SWD_detection, written in Matlab R2018b (MathWorks)) was used to automatically detect the SWD onset in real time and send a TTL trigger signal (via NI USB-6216, National Instruments) to control the DBS stimulator (STG 4002, Multichannel Systems). Videos were simultaneously recorded to monitor the locomotor activity. The EEG signal was saved for post hoc analysis. Video recording was synchronized to the EEG data via a LED light.

### *ex vivo* electrophysiology

For patch clamp recording, acute brain slices were prepared as described previously^52,55^. Briefly, *BK^WT/WT^* and *BK^DG/DG^*mice (from 3 weeks to 6 months) were anesthetized and decapitated. The MLT was processed into 300 µm coronal sections in ice-cold NMDG aCSF containing (in mM): 92 NMDG, 2.5 KCl, 1.2 NaH_2_PO_4_, 30 NaHCO_3_, 20 HEPES, 25 glucose, 5 sodium ascorbate, 2 thiourea, 3 sodium pyruvate, 10 MgSO_4_·7H_2_O, 0.5 CaCl_2_·2H_2_O (Titrated pH to 7.3-7.4 using concentrated HCl). The slices were then incubated in HEPES recovering solution (in mM): 92 NaCl, 2.5 KCl, 1.2 NaH_2_PO_4_, 25 NaHCO_3_, 20 HEPES, 25 glucose, 5 sodium ascorbate, 2 thiourea, 3 sodium pyruvate, 2 MgSO_4_·7H_2_O, 2 CaCl_2_·2H_2_O) for 60-min at room temperature. After incubation, the slices were transferred to a recording chamber and perfused (3 mL min^-1^) with artificial cerebrospinal fluid (aCSF) at 33 LJ. (in mM): 124 NaCl, 2.5 KCl, 1.2

NaH_2_PO_4_, 24 NaHCO_3_, 5 HEPES, 12.5 glucose, 2 MgSO_4_·7H_2_O, 2 CaCl_2_·2H_2_O. All solutions used for electrophysiology were equilibrated with 95% O_2_/5% CO_2_. Whole-cell recordings of MLT neurons were performed with a MultiClamp 700B amplifier and sampled at 10 kHz using a Digidata1550A A/D converter. All data acquisition and analyses were performed using the software pClamp 10.7 (Molecular Devices).

For action potential recording, the pipettes were filled with an intracellular solution containing the following (in mM): 125 K-gluconate, 15 KCl, 10 HEPES, 2 Mg-ATP, 0.3 Na-GTP, 10 disodium phosphocreatine, and 0.2 EGTA, adjusted to pH 7.25 with KOH. Pipette resistance was 3-7 MΩ. After GΩ-seal and membrane break-in, the membrane resting potential was monitored for 10 min until it is stabilized before recording of neuronal firing. For the rebound burst firing characterization in the *BK^WT/WT^* and *BK^DG/DG^* mice, the MLT neurons was held in current clamp at potential −70 mV through constant current injection. Rebound burst firing was induced by a 1 s, −200 pA current step. The number of the rebound bursts and inter-spike-interval (ISI) of the first and second rebound burst action potential (AP) were analyzed by the build-in function ‘Threshold Event Detection’ of the Clampfit 10.7 (Molecular Devices). AP90% duration was defined by action potential duration of 90% repolarization. The fAHP size was measured as the difference between the spike threshold and voltage minimum after the action potential. The AP90 and AHP were measured at the first action potential of each trace.

For optogenetic manipulation of MLT firing, similar as above, after the AAV2/9-CAG-ChR2-mCherry virus was expressed for 3 weeks, and the 300 µm coronal sections contained the MLT were prepared. To study the firing pattern of MLT neurons, a 20 Hz 5-ms blue laser light (473 nm, Inper Inc., Hangzhou, China) was delivered to the MLT neurons, which were held at −60 mV or −85 mV.

### Statistics

All the statistical analyses were performed using GraphPad Prism (GraphPad Software). Sample number (n) values are indicated in the results section and figure legends. All data are presented as the mean ± standard error of the mean (s.e.m.). Sample sizes were chosen based on standards in the field as well as previous experience with phenotype comparisons. No statistical methods were used to predetermine sample size.

## Data availability

All relevant data supporting the present study are available from the lead corresponding author upon reasonable request.

## Code availability

All relevant codes supporting the present study are available at GitHub https://github.com/superdongping/SWD_detection https://github.com/superdongping/c_fos_count

## Author contributions

H.Y. conceived and supervised the project, H.Y., P.D., H.Y., and W.M.G. designed the research. P.D. performed immunostaining, *ex vivo* electrophysiology, optogenetics and video-EEG. P.D. and K.B. conducted *in vivo* single-unit recording. P.D. and Y.L. designed the closed-loop SWD detection system and performed DBS. P.D., K.B., Y.L. wrote the MATLAB code and did data analysis. P.D. and H.Y. wrote the manuscript with input from all authors.

## Supporting information

Movie S1. Optogenetic stimulation of the midline thalamus suppressed absence seizures, promoted locomotion, and increased vigilance in BK-D434G mice.

Movie S2. Closed-loop deep brain stimulation (DBS) of the midline thalamus (MLT) suppresses absence seizures, promotes locomotion, and increases vigil

## Acknowledgement

We would like to thank Drs. Ru-Rong Ji, William Wetsel, James O’Connell McNamara, and Yang Zhang, for their comments and help on the manuscript. This work was supported by the Duke Institute for Brain Sciences (to H.Y.), the Duke University School of Medicine Bridge fund (to H.Y.) and American Epilepsy Society Post-Doctoral Fellowship 693905 (to P.D.).

## Supplementary figures

**Fig. S1.**
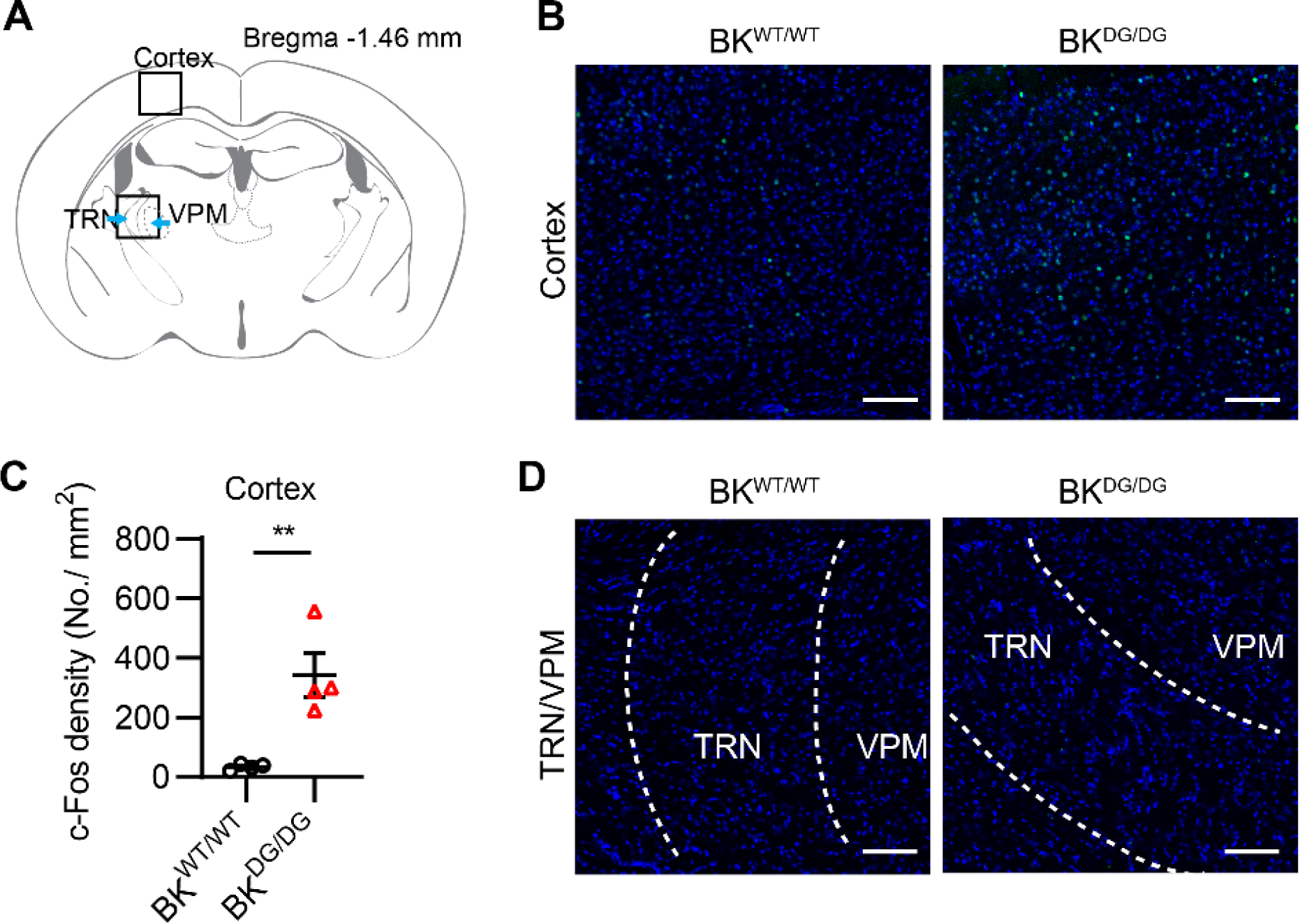
c-Fos expression is increased in the cortical pyramidal neurons but not the thalamic reticular nucleus (TRN) or posteromedial nucleus (VPM) from BK-D434G mice. (**A**) A coronal atlas showing the brain regions in panels B and D. The blue arrows indicate the position of the TRN and the VPM. (**B**) Representative images of c-Fos expression in the cortex of the *BK^WT/WT^* and *BK^DG/DG^*mice. (**C**) Statistics of c-Fos density of BK-D434G mice in the cortex. Two-tailed unpaired Student’s *t*-test, t_6_ = 4.182, *P* = 0.0058. (**D**) Representative images of c-Fos expression in the TRN and the VPM of the *BK^WT/WT^*and *BK^DG/DG^* mice. Scale bar 100 μm.

**Fig. S2.**
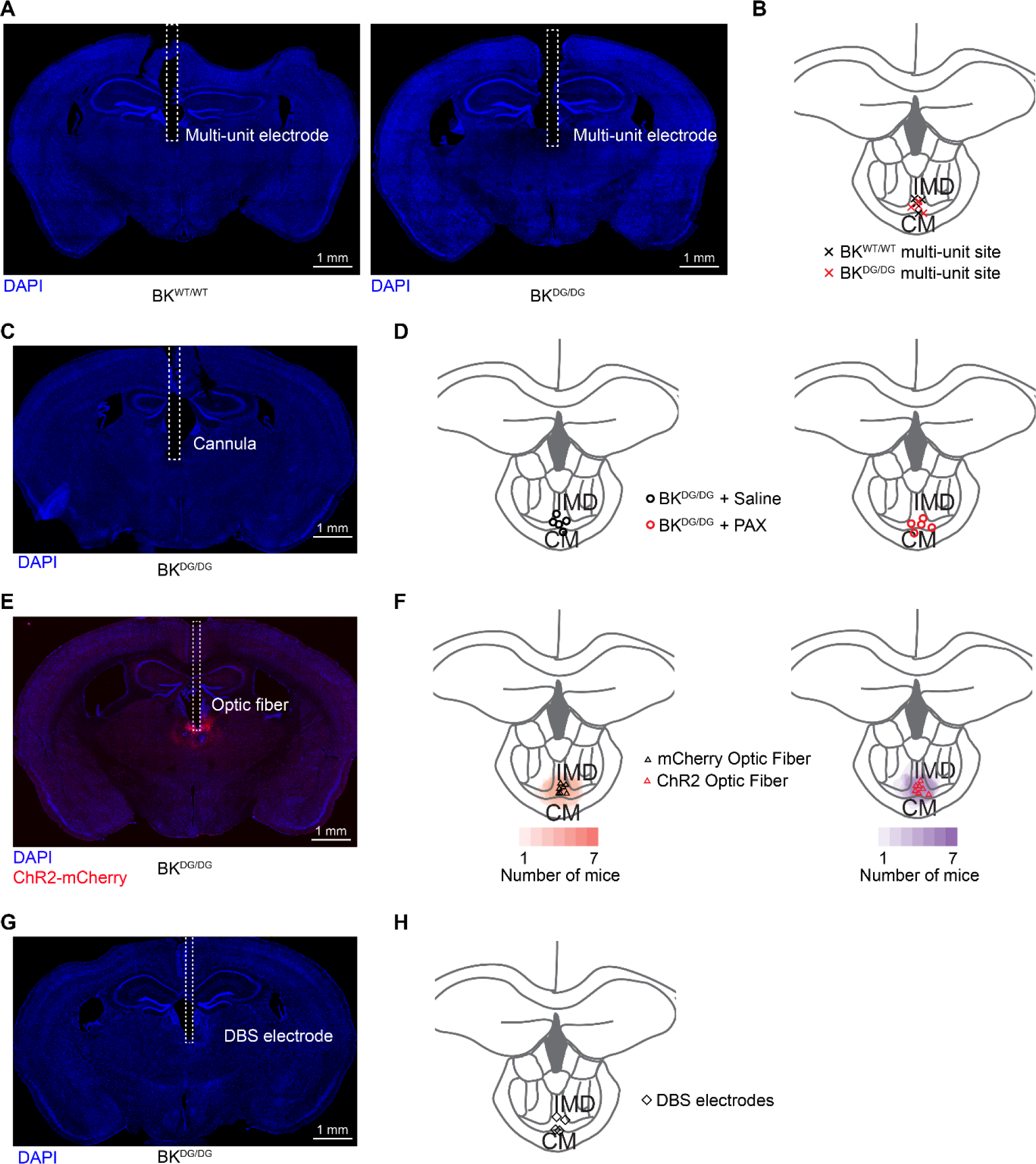
Histological verification of the placement of single unit multi-electrode array, cannula, optogenetic fiber, and DBS electrode array in the MLT. (**A**) Representative brain images from a wildtype (BK^WT/WT^) mouse and a D434G mutant (BK^DG/DG^) mouse with an electrode array implanted in the MLT. (**B**) Summary of the positions of single-unit electrode tips from all mice. (**C**) Representative brain images a BK^DG/DG^ mouse with a cannula implanted in the MLT. (**D**) Summary of the positions of cannula tips from all mice injected with saline (left) or PAX (right). **(E)** Representative brain images from a mouse with viral ChR2 or mCherry expression (red) and an optical fiber (dotted lines) implantation in the MLT of a BK^DG/DG^ mouse. **(F)** Summary of the sites for viral expression and optical fiber implantation (Left: mCherry; Right: ChR2). (**G**) Representative brain images a BK^DG/DG^ mouse with DBS electrodes implanted in the MLT. (**H**) Summary of the positions of DBS electrodes sites from all mice.

**Fig. S3.**
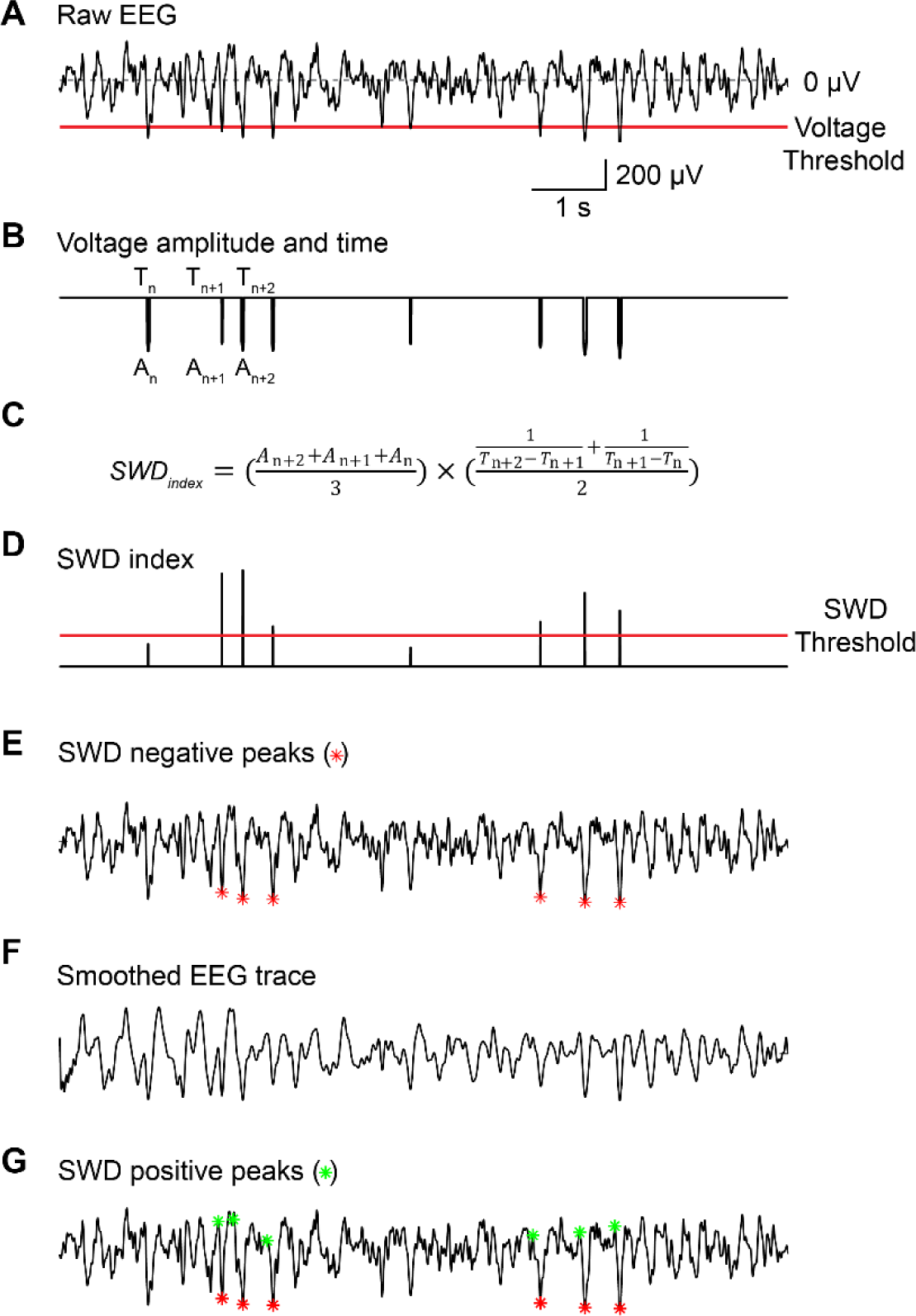
The procedure to automatically detect the timestamps of spike-and-wave-discharges (SWDs). (**A**) Representative EEG raw trace from a BK^DG/DG^ mouse. A voltage threshold is empirically set to −300 μV to facilitate detecting SWD peaks. (**B**) The raw EEG is filtered/transformed based on the voltage threshold. The voltage amplitude beyond the threshold and the corresponding time are extracted during this process. (**C**) The equation used to calculate the weighted SWD index (SWD_index_), which equals the multiplication of the average of the voltage amplitudes (A) from three consecutive events and the average of the frequencies of these events (see Methods for details). (**D**) SWD indexes are calculated and plotted. To minimize false positive SWD detection, only the top 75% of SWD_index_ values are included from further analysis, and the corresponding EEG signal is considered as an SWD event. (**E**) The negative peaks of the SWD events in the raw EEG trace are labeled with red asterisks. (**F**) The raw EEG trace is subsequently smoothened prior to the detection of the positive peaks of SWD events. (**G)** The positive peaks of SWD events in the raw EEG trace are labeled with green asterisks.

**Fig. S4.**
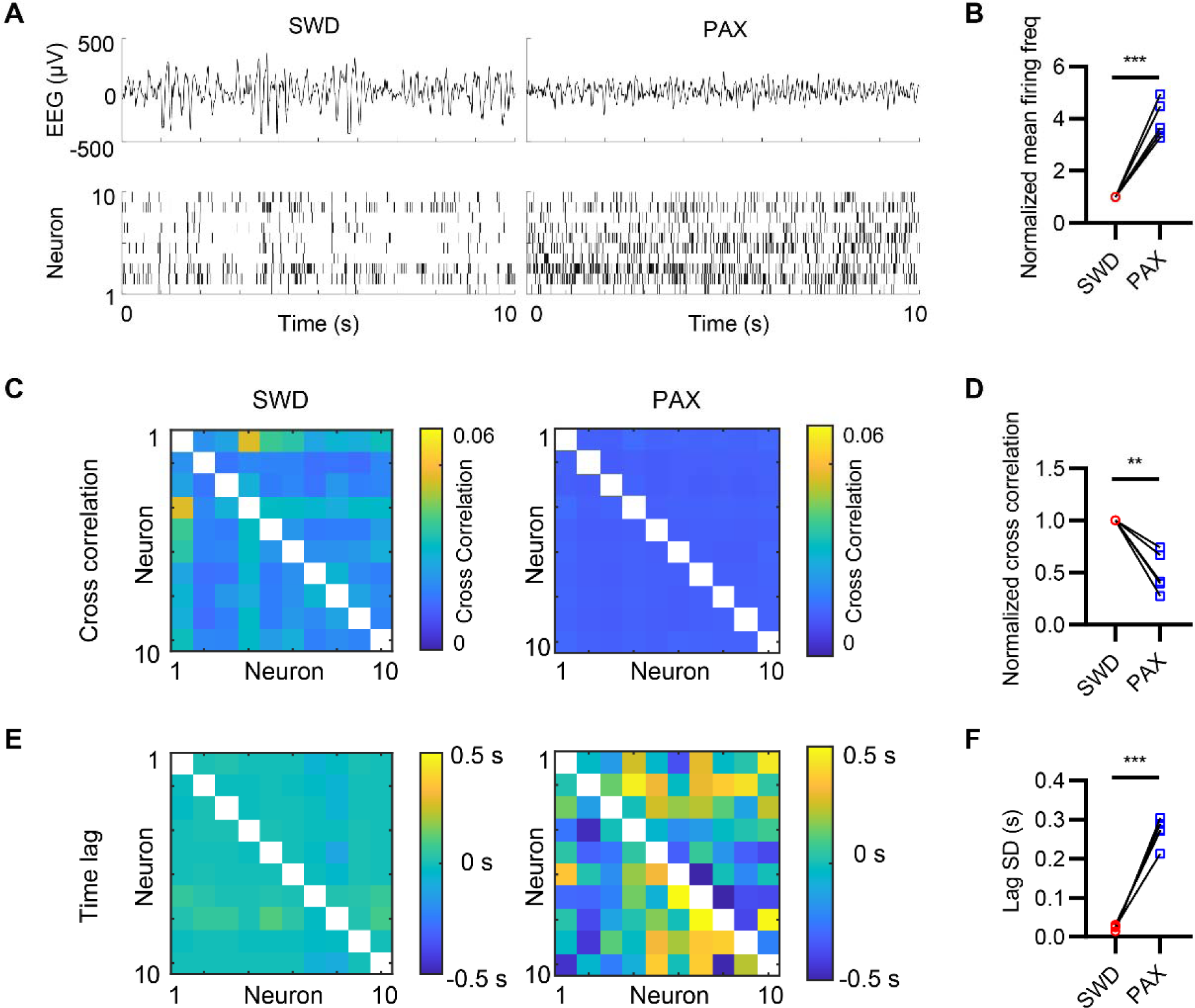
Inhibiting BK channels with paxilline (PAX) desynchronizes MLT firing *in vivo*. (**A**) Representative raw EEG traces (top) and single-unit signals from different BK^DG/DG^ MLT neurons (bottom) during the ictal (SWD) period before PAX administration and a period after PAX administration (PAX). (**B**) Relative change of mean firing frequency normalized to the SWD period. Two-tailed paired Student’s *t*-test, t_4_ = 9.297, *P* = 0.0007. (**C**) Cross correlation indexes among different MLT neurons during SWD and PAX periods. (**D**) Relative cross correlation indexes normalized to the SWD period. Two-tailed paired Student’s *t*-test, t_4_ = 5.688, *P* = 0.0047. (**E**) Mean time lag among different MLT neurons during SWD and PAX periods. (**F**) Standard deviation (SD) of time lag among different neurons during SWD and PAX periods. Two-tailed paired Student’s *t*-test, t_4_ = 14.41, *P* = 0.0001. n = 5 mice. In all plots and statistical tests, summary graphs show mean ± s.e.m.

**Fig. S5.**
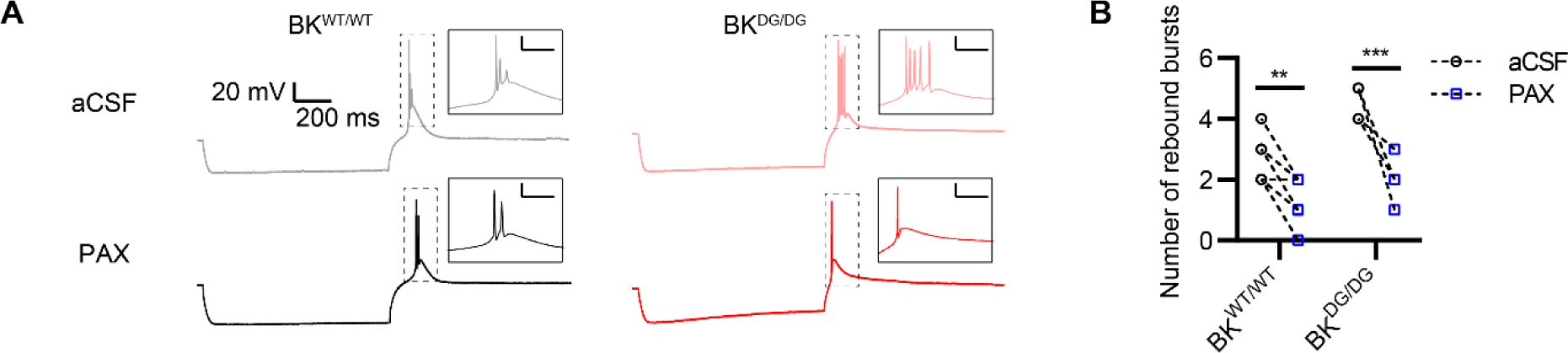
Pharmacological inhibition of BK channels suppresses burst firing in MLT neurons. (**A**) Representative low-threshold rebound burst firing from the MLT neurons from BK^WT/WT^ (black trace) or BK^DG/DG^ (red trace) mice before (top, aCSF) and after (bottom) application of 10 µM PAX. The rebound burst firing was elicited by a −200-pA negative current injection for 1 s. (**B**) Number of the rebound burst firing number from the MLT of BK^WT/WT^ and BK^DG/DG^ neurons before and after application of 10 µM PAX. Two-way ANOVA, *F*_(1,12)_ = 44.28, *P* < 0.0001. n = 10 neurons from three BK^WT/WT^ mice, n = 4 neurons from two BK^DG/DG^ mice.

**Fig. S6.**
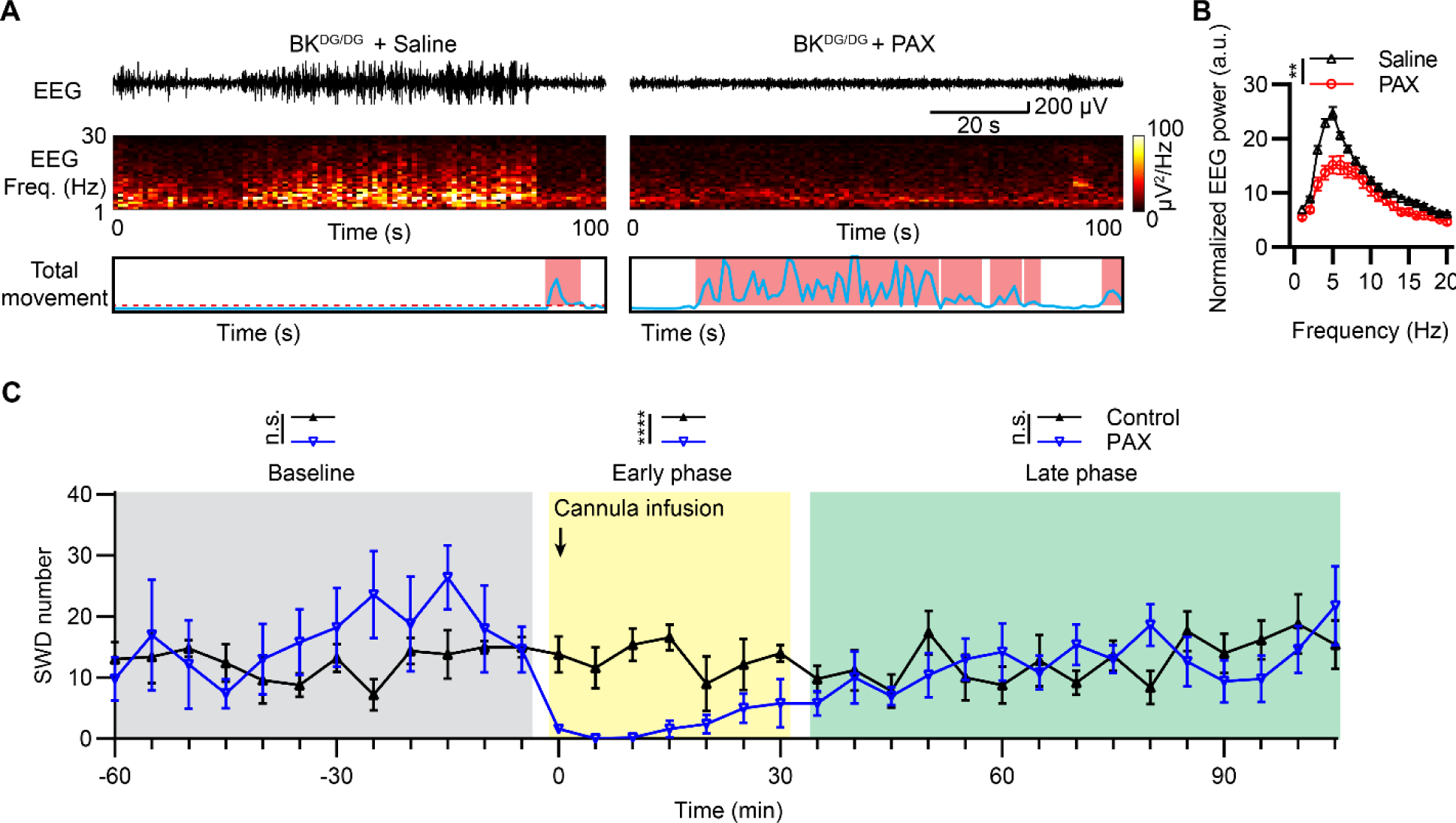
Local inhibition of BK channels in the MLT suppresses absence seizures. (**A**) Representative EEG traces (top), corresponding spectrograms (middle), and total movement (bottom) from the *BK^DG/DG^* mice administrated with saline (left panel) or 1 µL, 1 µM PAX (right panel) via cannula. (**B**) Summary of power spectral density of EEG recorded from *BK^DG/DG^* mice administrated with saline or PAX. Two-way ANOVA, *F*_(1,10)_ = 14.60, *P* = 0.0034. n = 6 mice per group. (**C**) Time course of the cannula administration of PAX (blue) and saline control (black) on the spontaneous SWDs of the *BK^DG/DG^* mice (bin size = 5 min). The drug effects were empirically divided into 4 different phases: baseline phase, 60 min prior to injection (grey box, two-way repeated-measures ANOVA, F_(1,8)_ = 0.2399, *P* = 0.6374); early phase, 30 min post injection (yellow box, two-way repeated-measures ANOVA, F_(1,8)_ = 50.61, *P* = 0.0001); and late phase, from 35 to 105 min post injection (green box, two-way repeated-measures ANOVA, F_(1,8)_ = 0.1402, *P* = 0.7178). n = 5 mice for each group.

**Fig. S7.**
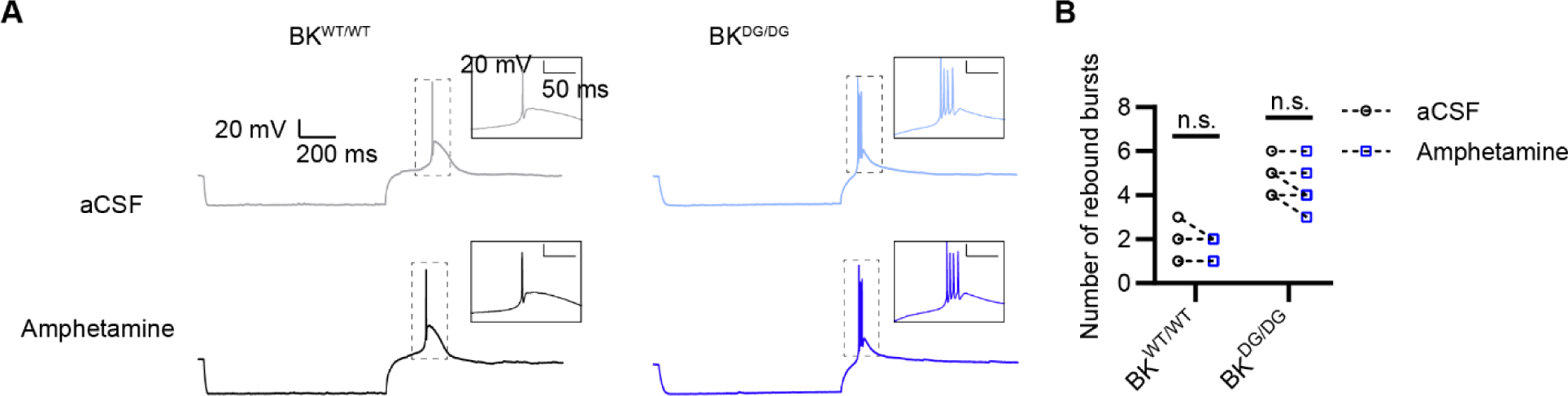
Amphetamine has a negligible effect on the rebound burst firing of the BK-D434G MLT neurons. (**A**) Representative rebound burst spikes of the MLT neurons from BK^WT/WT^ (black trace) or BK^DG/DG^ (blue trace) neurons before (top) and after (bottom) applying 10 µM D-amphetamine sulfate. The rebound burst firing is elicited by a −200-pA negative current injection for 1 s. (**B**) Quantification of the rebound burst spike numbers from BK^WT/WT^ and BK^DG/DG^ MLT neurons before and after application of amphetamine. Two-way ANOVA, *F*_(1,10)_ = 3.462, *P* = 0.0924. n = 6 neurons from three BK^WT/WT^ mice, n = 6 neurons from three BK^DG/DG^ mice.

## Captions for Movies

**Movie S1. Optogenetic stimulation of the midline thalamus suppressed absence seizures, promoted locomotion, and increased vigilance in BK-D434G mice.**

This video shows a simultaneous video-EEG recording of a freely moving BK-D434G mouse, subjected to either no optogenetic stimulation, single optogenetic stimulation (3 seconds, 20 Hz, 5 ms pulse width), or continuous stimulation (10 minutes, 20 Hz, 5 ms pulse width). The mouse experienced frequent spike-and-wave discharges (SWDs) and behavioral arrest without optogenetic stimulation. Optogenetic stimulation suppressed SWDs and promoted vigilance as evidenced by increased locomotor activities such as exploration, walking, and turning.

**Movie S2. Closed-loop deep brain stimulation (DBS) of the midline thalamus (MLT) suppresses absence seizures, promotes locomotion, and increases vigilance in BK-D434G mice.**

This video shows a representative simultaneous video-EEG recording of a freely moving BK-D434G mouse when the closed-loop DBS is either on or off. A green LED indicates the detection of SWDs from the EEG signal (for more details, refer to Fig 6A, Fig. S3, and the Methods section). For the “DBS on” trial, a DBS stimulation (5 seconds, 20 Hz, 1 ms pulse width, 50 µA) was delivered to the MLT of the mouse immediately after detecting an SWD event. The DBS suppressed SWDs and promoted vigilance as evidenced by increased locomotor activities such as exploration, walking, and turning. In contrast, during the ‘DBS off’ trial, SWDs were detected but no DBS stimulation was delivered to the MLT. The mouse continued to undergo absence seizure attacks as evidenced by behavioral arrest and frequent SWDs.

